# Aberrant cell state plasticity mediated by developmental reprogramming precedes colorectal cancer initiation

**DOI:** 10.1101/2022.09.22.509032

**Authors:** Pratyusha Bala, Jonathan P. Rennhack, Daulet Aitymbayev, Clare Morris, Sydney M. Moyer, Gina N. Duronio, Paul Doan, Zhixin Li, Xiaoyan Liang, Jason L. Hornick, Matthew B. Yurgelun, William C. Hahn, Nilay S. Sethi

## Abstract

Cell state (phenotypic) plasticity is a carefully regulated feature of adult epithelial cells that enables adaptive responses to injury, inflammation, and other forms of stress. Aberrant expansion of the normally restricted capability for cell state plasticity to escape terminal differentiation is a critical aspect of neoplasia. The nongenetic factors and specific programs that mediate aberrant cell state plasticity and impaired differentiation require deeper characterization to understand this elusive aspect of cancer pathogenesis. Using genetically engineered and carcinogen-induced murine models of intestinal neoplasia, we demonstrate that impaired differentiation is a conserved event preceding cancer development. Single cell RNA-sequencing (scRNA-seq) of neoplastic intestinal lesions from both mouse models and a patient with familial adenomatous polyposis revealed that cancer initiates by adopting an aberrant transcriptional state characterized by nonoverlapping expression of a regenerative pathway, marked by *Ly6a* (Sca-1), and a fetal intestinal program, positive for *Tacstd2* (Trop2). Genetic inactivation of *Sox9* prevented adenoma formation in Apc^KO^ mice, obstructed emergence of aberrant regenerative and fetal intestinal programs, and restored multi-lineage differentiation by scRNA-seq. Expanded chromatin accessibility at regeneration and fetal genes upon Apc inactivation was reduced by concomitant Sox9 suppression. These studies indicate that aberrant cell state plasticity mediated by unabated regenerative activity and developmental reprogramming precedes cancer development.

## INTRODUCTION

Cell state plasticity – the ability of cells to acquire new states via differentiation programs - is a critical cellular feature of embryogenesis, allowing for tissue specification during development(*1*), and adult homeostasis, enabling tissue repair and regeneration (*2*). While cellular plasticity progressively decreases during development, it remains a normal albeit restricted feature of adult epithelial cells that facilitates adaptation to injury, inflammation, among other forms of stress. Eukaryotes rely on epigenetic governance among other layers of regulation to endow cell state plasticity while avoiding properties that permit neoplasia(*3*). Recent studies are providing key evidence that unrestricted cellular plasticity underlies neoplastic initiation, molecular heterogeneity and suboptimal response to therapy(*4*–*8*). Impaired differentiation is a key mechanism by which neoplastic cells unlock cell state plasticity(*6*, *9*). While differentiation blocks are a well-studied aspect of premalignant hematological conditions and leukemogenesis, its significance in colorectal cancer (CRC) is lesser appreciated.

The rapidly renewing adult intestinal epithelium engenders inherent cellular plasticity. Crypt-restricted Lgr5^+^ intestinal stem cells (ISC) and their immediate progeny replicate frequently to spawn a steady stream of postmitotic differentiated cells that function in barrier protection, absorption, and hormone secretion in the villus before dying(*10*). The key pathways that regulate renewal and differentiation include wingless/integrated (WNT) and bone morphogenetic protein (BMP) signaling cascades, respectively. WNT pathway activity maintains stem cell reservoirs and crypt homeostasis whereas BMP signaling supports differentiation of progenitors into mature enterocytes, establishing a crypt-villus gradient(*11*, *12*) (Fig. 1A). There are limited differences in chromatin accessibility and regulatory histone marks between ISCs and their differentiated descendants despite distinct gene expression profiles(*13*, *14*), underscoring the intrinsic plasticity among cell types that facilities tissue repair and regeneration. While disruption of stem cell and differentiation programs is observed in CRC, which is the third most common and second most deadly malignancy worldwide (*15*), we have yet to define regulators of cell state plasticity in cancer initiation.

**Figure 1.**
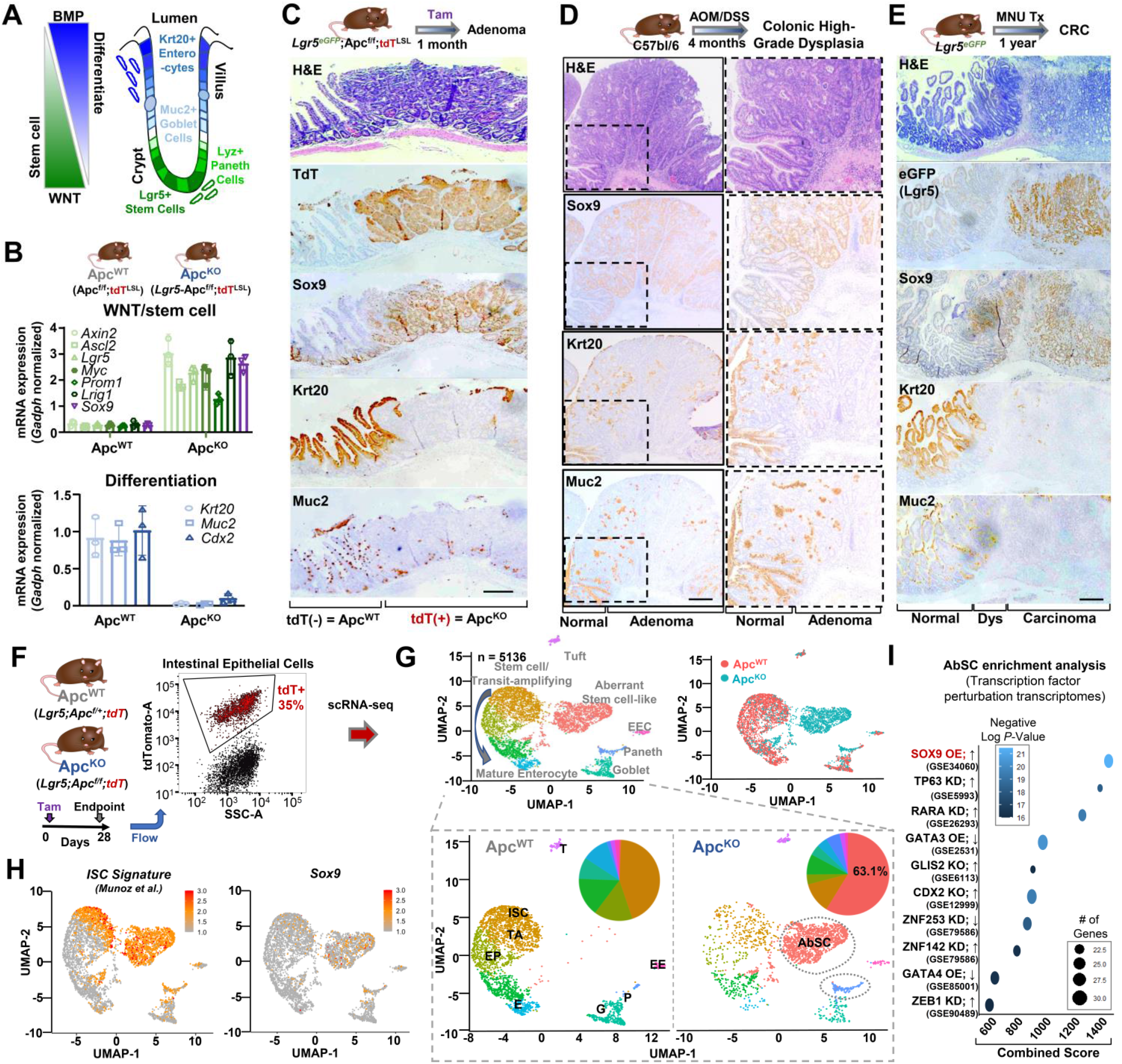
Impaired differentiation is a conserved mechanism of aberrant cell state plasticity in mouse models of intestinal neoplasia. A. Schematic depicting WNT and BMP signaling gradient in the intestines. Stem cells and Paneth cells reside in the crypt whereas differentiated cell types are found in the villus. B. Relative mRNA expression of Wnt/stem cell (top) and differentiation genes (bottom) in intestinal tissue derived from indicated mice by qRT-PCR; mean ± S.D. of three biological replicates. C. Representative images of intestinal lesions from *Lgr5^Cre^;Apc^f/f^;R26^tdT^* mice including hematoxylin and eosin (H&E) staining and tdTomato (tdT), Sox9, Krt20, and Muc2 immunohistochemistry (IHC); scale bar = 250 μM. D. Representative images of intestinal lesions from AOM/DSS treated mice including H&E staining Sox9, Krt20, and Muc2 IHC. Normal and adenoma regions labeled; scale bar = 250 μM. E. Representative images of intestinal lesions from MNU treated Lgr5^eGFP^ mice including H&E staining and eGFP (Lgr5 expressing cells), Sox9, Krt20, and Muc2 IHC. Normal, dysplastic and carcinoma regions labeled. Scale bar = 250 μM. F. tdT+ intestinal epithelial cells isolated by fluorescence-activated cell sorting (FACS) from *Lgr5^Cre^; Apc^f/+^;R26^tdT^* (control) and *Lgr5^Cre^;Apc^f/f^;R26^tdT^* (experimental) mice 28 days following tamoxifen induction. G. UMAP representation of single cell transcriptome profiling of tdT+ epithelial cells from *Lgr5^Cre^; Apc^f/+^;R26^tdT^* (control) and *Lgr5^Cre^;Apc^f/f^;R26^tdT^* (experimental) mice colored by cell type (top left panel) and sample identity (top right panel). UMAP of separated samples along with pie chart indicating distribution of cell types (bottom panel). ISC = stem cell, TA = transit amplifying, EP = enterocyte progenitor, E = mature enterocyte, G = goblet cell, P = Paneth cell, EE = enteroendocrine, T = tuft cell, AbSC = aberrant stem cell-like. H. UMAP representation of intestinal stem cell signature (Munoz et al.) (left) Sox9 (right). I. Transcription factor perturbation gene ontology analysis (EnrichR) of top 100 genes upregulated in AbSC cluster.

The majority of sporadic CRCs initiate as a premalignant adenoma harboring genomic alterations that constitutively activate WNT signaling(*16*, *17*), most often through loss-of-function *APC* mutations(*18*, *19*). A key component of a cytoplasmic destruction complex, APC restricts WNT signaling by facilitating ubiquitin-mediated degradation of β-catenin(*20*–*22*). Once APC function is compromised, β-catenin accumulates in the nucleus, where it can operate as the primary effector of aberrant WNT signaling(*23*, *24*). ISC expansion and imbalanced crypt-villus homeostasis are known outcomes of *Apc* deletion and constitutive β-catenin activity in mouse models of intestinal neoplasia(*25*, *26*). Beyond these cellular outcomes, the nongenetic transcriptional consequences and cell state heterogeneity following deleterious *APC* mutations are poorly understood.

Restoring expression of functional APC can reinstate WNT pathway regulation and suppress cancer initiation(*27*–*29*). Inhibiting WNT-dependent Lgr5^+^ stem cells can impair tumor growth and metastasis (*30*, *31*). Translating these concepts into effective therapy, however, has proven difficult as WNT pathway inhibitors have not advanced in preclinical and clinical testing(*32*). By defining key mediators that facilitate neoplastic initiation, we may uncover new therapeutic strategies for CRC. Here, we characterize critical early molecular events of intestinal neoplasia using histopathological analyses, single cell transcriptomics, chromatin accessibility assays, and organoid experiments in distinct mouse models and human specimens, uncovering developmental reprogramming as an important early step leading to aberrant cell state plasticity preceding cancer formation.

## RESULTS

### Impaired differentiation is a conserved event in multiple mouse models of intestinal neoplasia

To study the molecular underpinnings of aberrant cell state plasticity, we employed two genetically engineered and two carcinogen-induced murine models of intestinal neoplasia. *Lgr5*^eGFP-IRES-CreERT2^;*Apc*^loxP-exon14-loxP^(*Lgr5^Cre^;Apc^f/f^*) genetically engineered mice develop hundreds of adenomas in the small intestines following tamoxifen-induced conditional deletion of Apc(*33*) in Lgr5^+^ intestinal stem cells (ISC)(*10*, *34*) (fig. S1, A and B). Compared to controls, adenomas from *Lgr5^Cre^;Apc^f/f^* mice displayed elevated mRNA expression of canonical WNT downstream targets (*Axin2, Myc*) and stem cell genes (*Ascl2, Lgr5, Prom1, Lrig1*), including *Sox9* (Fig. 1B), consistent with known biology(*35*, *36*). In contrast, markers of differentiated intestinal cells (e.g. *Cdx2*) of both absorptive (e.g. *Krt20*) and secretory lineages (e.g. *Muc2*)(*37*) were downregulated in *Lgr5;Apc^f/f^* intestinal adenomas (Fig. 1B). To validate these findings, we examined another previously published genetically engineered mouse model in which inducible Apc knockdown (KD) with or without mutant K-ras^G12D^ activation in Lgr5^+^ cells displayed robust formation of colonic adenomas(*27*). In agreement with the *Lgr5^Cre^;Apc^f/f^* model, genes associated with stem cell activity were upregulated whereas intestinal differentiation genes were suppressed in Apc KD colonic adenomas by bulk RNA-seq (fig. S1C). We also analyzed protein expression of Sox9 in early intestinal adenomas from *Lgr5^Cre^;Apc^f/f^* mice that were crossed with *R26^LSL-tdTomato^* (*R26^tdT^* or *tdT^LSL^*) mice to label Apc^KO^ cells. Compared to adjacent normal intestinal epithelium in which Sox9 expression is restricted to the crypt base, tdT^+^ Apc^KO^ lesions demonstrated ectopic and elevated expression of Sox9 throughout the lesion, which coincided with absence of differentiation markers Krt20 and Muc2 (Fig. 1C). These observations suggest that Apc inactivation is sufficient to impair intestinal differentiation and leads to inappropriate Sox9 expression in adenomas.

We next evaluated two mouse models of carcinogen-induced small intestinal and colonic neoplastic lesions to ask whether Sox9 activation and impaired differentiation are universal events in cancer initiation that can be achieved by random mutagenesis *in vivo*. The first, well-established model consists of exposing mice to the procarcinogen azoxymethane (AOM) and irritant dextran sodium sulfate (DSS) (methods), which leads to colonic lesions ranging from low-grade dysplasia to intramucosal adenocarcinoma after several months. Consistent with our observations in the *Lgr5^Cre^;Apc^f/f^;R26^tdT^* model, high-grade dysplastic colonic lesions demonstrated robust Sox9 overexpression and loss of Krt20^+^ enterocytes and Muc2^+^ goblet cells (Fig. 1D and fig. S1D). In the second model, mice were exposed to drinking water containing N-methyl-N-nitrosourea (MNU), a potent carcinogen with gastrointestinal tropism, and a subset developed poorly differentiated intestinal carcinomas after one year. *Lgr5^Cre^* mice treated with MNU showed marked upregulation of Sox9 in premalignant and adenocarcinoma lesions (Fig. 1E and fig. S1E), consistent with an early role in cancer development. Differentiation markers Krt20 and Muc2 were absent in premalignant and adenocarcinoma lesions. Elevated expression of Lgr5 as indicated by eGFP immunohistochemistry (IHC) was found in intestinal adenocarcinomas but not their associated premalignant lesions, underscoring the central importance of stem cell activation in progression to malignancy (Fig. 1E). These results implicate impaired differentiation and Sox9 activation as a common pathway to CRC initiation.

To characterize the transcriptional program enabling impaired differentiation, we performed single cell RNA-sequencing (scRNA-seq) on the *Lgr5^Cre^;Apc^f/f^;R26^tdT^* genetic model. tdT^+^ epithelial cells were isolated from *Lgr5^Cre^;Apc^f/+^;R26^tdT^* (controls that are phenotypically wildtype) and *Lgr5^Cre^;Apc^f/f^;R26^tdT^* mice by fluorescence-activated cell sorting (FACS) 28 days following tamoxifen induction (Fig. 1F). Unsupervised clustering followed by uniform manifold approximation and projection (UMAP) representation of single-cell gene expression profiles revealed that control Apc^WT^ cells displayed the expected distribution of cell lineages based on intestinal markers(*38*) (fig. S2, A to C). In contrast, Apc inactivation led to a 4-fold enrichment in Paneth cells (5.9% vs 1.5%) and the emergence of cells that exhibited a new transcriptional state (63.1% vs 0%) at the expense of normal ISCs and differentiated cell types, including absorptive progenitors, mature enterocytes, goblet cells, and, to a lesser extent, enteroendocrine cells (Fig. 1G). Indeed, tdT^+^ Apc^KO^ intestinal lesions demonstrated a greater number of Paneth cells by Lyz IHC (fig. S3A), which can also be found at earlier timepoints of Apc inactivation(*35*, *39*). Apc deletion led to a greater percentage of Lgr5^+^ cells as indicated by eGFP FACS (fig. S2D), suggesting that ISC activity is upregulated in Apc^KO^ cells, agreeing with a vast literature(*40*) and supported by the enrichment of published ISC signatures (table S1)(*38*, *41*) (Fig. 1H and fig. S2A). A deeper evaluation revealed that select canonical ISC markers were not consistently found in Apc^KO^ cells, including *Olfm4* (fig. S3, B and C). In contrast, Apc^KO^ cells expressed higher levels of *Sox9* compared to normal ISC (Fig. 1H), which was supported by universal Sox9 expression in tdT^+^ intestinal lesions from *Lgr5^Cre^;Apc^f/f^;R26^tdT^* mice (Fig. 1C). Furthermore, evaluating transcription factor perturbation gene-sets that were enriched among upregulated genes in Apc^KO^ cells (table S1) revealed that a Sox9 overexpression transcriptional program scored the strongest (Fig. 1I, adjusted p-value = 3.26 x 10^-22^, table S2). These observations suggest that Apc deletion leads to a new transcriptional state with selective ISC transcriptional activity.

### Reactivation of genes associated with fetal intestines upon Apc loss

Despite showing an enrichment of most canonical ISC genes, the transcriptional profile of Apc^KO^ cells were found in one distinct cluster (Fig. 1G), expressing genes not found in normal intestinal cells; we therefore refer to these cells as aberrant stem cell-like (AbSC). To characterize the AbSC transcriptional program in greater depth, searching for features that explain its distinguished gene expression profile, we examined the differentially expressed genes in the AbSC cluster relative to all other intestinal clusters. Consistent with our previous results, genes associated with differentiatied enterocytes (blue) were downregulated whereas ISC markers and WNT pathway targets (green) were upregulated (Fig. 2A). Genes associated with interferon (IFN) signaling (fig. S4, A and B) were significantly upregulated in AbSCs, which matched gene ontology analyses implicating IFN pathways and transcription factors (e.g., RELA, NF-κB) (fig. S4C and table S2). Genes associated with regeneration (pink) and fetal intestines (purple) were also uniquely and significantly upregulated in the AbSC cluster (Fig. 2A). Indeed, a regeneration gene expression signature derived from an experimental intestinal Helminth infection model (Nusse, table S1)(*42*) was specifically upregulated in AbSCs (Fig. 2B). Notably, this gene expression signature is also associated with IFN-γ signaling, indicating a potential link between regeneration and IFN-γ signaling(*42*).

**Figure 2.**
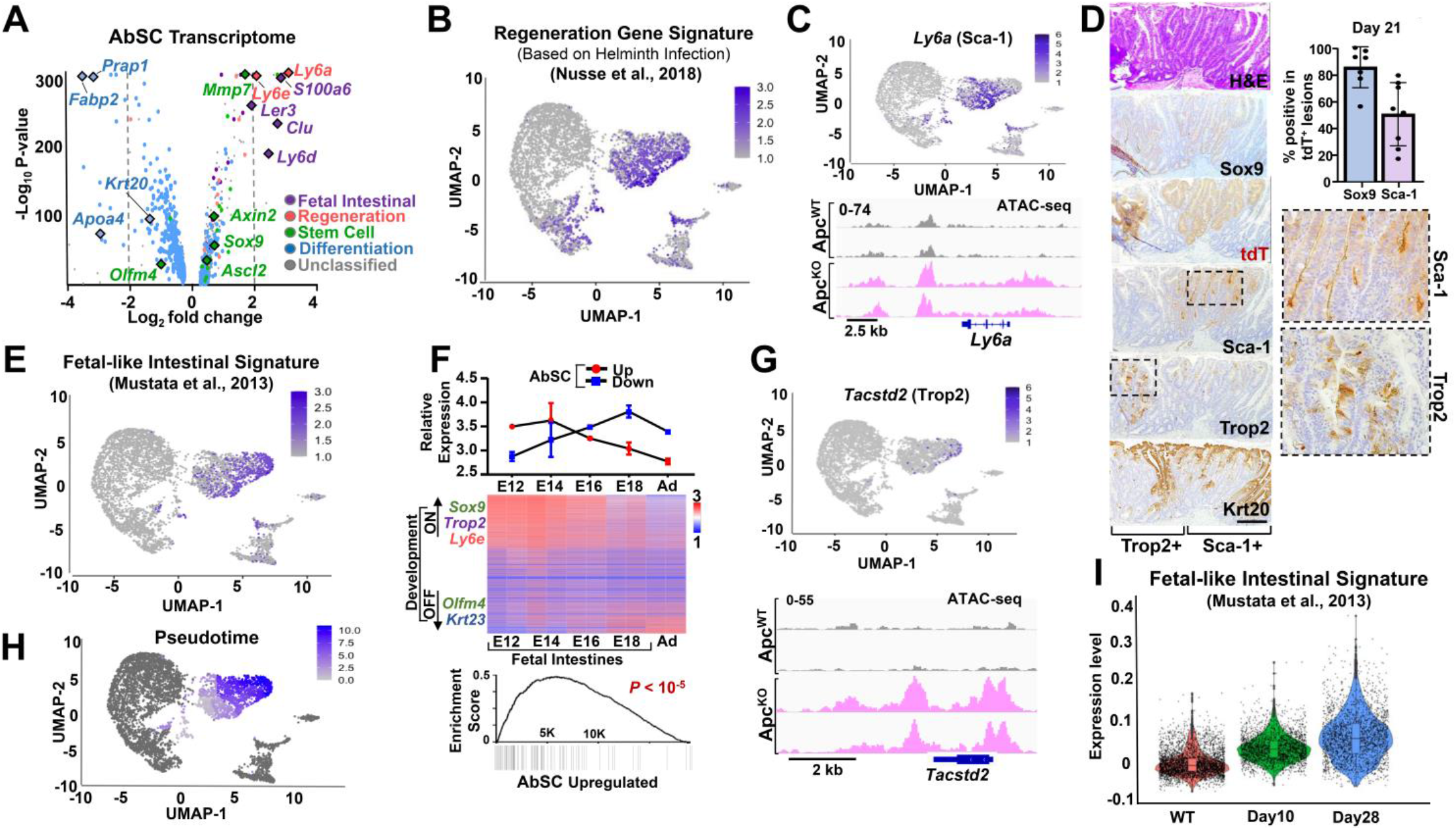
Reactivation of genes associated with fetal intestinal development upon *Apc* inactivation. A. Volcano plot showing differentially expressed genes from AbSC cluster. Enterocyte (blue), stem cell (green), regeneration (salmon), and fetal intestines (purple) genes are highlighted. B. Normalized regeneration gene signature expression (Nusse et al., 2018) on UMAP plot C. Normalized *Ly6a* expression on UMAP plot (top). Chromatin accessibility at *Ly6a* genomic locus in tdT^+^ cells isolated by FACS from *Lgr5-tdT* and *Lgr5*-Apc^f/f^-tdT mice by ATAC sequencing (bottom). D. Representative images of intestinal lesions from *Lgr5*-Apc^f/f^-tdT mice including hematoxylin and eosin (H&E) staining and tdTomato (tdT), Sca-1 (quantification on top right), Trop2, and Krt20 immunohistochemistry (IHC); scale bar = 250 μM. E. Normalized fetal-like intestinal gene signature expression (Mustata et al., 2013) on UMAP plot F. Average expression of differentially upregulated (red) and downregulated (blue) genes across fetal intestines at indicated timepoints of mouse development by RNA-seq (Banerjee et al.). Heatmap representing differentially expressed genes in E12 fetal intestines compared to adult intestines ranked by fold change; a normalized enrichment score of upregulated genes in AbSC is shown on the right (p < 1×10^-5^). E = embryonic, number indicates day, Ad = adult G. Normalized *Tacstd2* expression on UMAP plot (top). Chromatin accessibility at *Tacstd2* genomic locus in tdT^+^ cells isolated by FACS from *Lgr5-tdT* and *Lgr5*-Apc^f/f^-tdT mice by ATAC sequencing (bottom). H. Pseudotime analysis of AbSCs using Monocle 3. I. Violin plot indicated expression of fetal-like intestinal gene signature expression (Mustata et al., 2013) in tdT+ intestinal cells from *Lgr5-tdT* control and *Lgr5*-Apc^f/f^-tdT at day 10 and 28 following tamoxifen induction. *P*-value calculated by Wilcoxon rank sum test with Bonferroni correction.

Among regeneration genes within AbSCs, *Ly6a* and *Ly6e* consistently showed the strongest and most abundant expression (Fig. 2C and fig. S4D). While Apc inactivation lead to minor overall chromatin accessibility changes at regeneration genes (fig. S4E), there was greater access at *Ly6a* and *Ly6e* genomic loci by assay for transposase accessible-chromatin (ATAC-seq)(*43*) (Fig. 2C and fig. S4D). We decided to focus on *Ly6a* (also referred to as stem cell antigen-1 [Sca-1]) as a marker for validation studies due to its (1) selective high-level expression in a substantial fraction of AbSC cells (63.1%), (2) absence from normal intestinal clusters, and (3) association with stem cell properties in the hematopoietic system (*44*–*46*). Greater chromatin accessibility at the *Ly6a* genomic locus upon Apc deletion corresponded to elevated *Ly6a* mRNA expression by RT-PCR (fig. S4F). *Lgr5^Cre^;Apc^f/f^;R26^tdT^* mice displayed selective Sca-1 expression in tdT^+^ intestinal lesions and, within these lesions, Sca-1 was expressed in 30-40% of tdT^+^ cells (Fig. 2D). Sca-1^+^ cells also expressed Sox9, which was found in approximately 85% of tdT^+^ cells (Fig. 2D), but did not consistently express Lgr5/eGFP) or Olfm4 (fig. S4, G and H). To examine self-renewal properties of Sca-1^+^ AbSCs, we performed organoid-forming assays using tdT^+^ cells isolated by FACS. Consistent with *in situ* obsevations, ~28% of tdT^+^ cells from *Lgr5^Cre^;Apc^f/f^;R26^tdT^* mice demonstrated Sca-1 protein expression by FACS, whereas tdT^+^ cells from *Lgr5-tdT* mice showed no expression (fig. S4I). Per 1000 cells plated, Sca-1^+^ cells showed a 7.4-fold greater organoid forming ability than Sca-1^-^ cells isolated from *Lgr5^Cre^;Apc^f/f^;R26^tdT^* mice (fig. S4, J and K). Dissociated Sca-1^+^ organoids passaged into secondary cultures maintained a 5.9-fold greater organoid forming ability than Sca-1^-^ cells (fig. S4L), suggesting population stability and preserved regenerative properties. These data suggest that sustained regenerative transcriptional activity is a feature of neoplastic initiation and contributes to self-renewal capacity.

Fetal intestinal genes were also upregulated in AbSCs (Fig. 2A). Consistently, two relatively nonoverlapping fetal-like gene signatures(*47*, *48*) (Mustata and Vallone, table S1) mapped specifically to the AbSC cluster (Fig. 2E and fig. S5, A and B). To confirm these associations, we also interrogated bulk gene expression data directly extracted from intestines at different stages of embryonic development(*1*). AbSC downregulated genes had stronger expression in late development (E18) and adult intestines compared to earlier stages of intestinal development (Fig. 2F, E12-E16). By contrast, AbSC upregulated genes are significantly enriched among genes preferentially expressed in embryonic relative to adult intestines (Fig. 2F), which include *Sox9, Tacstd2* (encoding Trop2), and *Ly6e*. Among fetal intestinal genes, *Tacstd2* showed the most specific expression in AbSCs, marking atleast 11% of the cluster (Fig. 2G), which was confirmed by Trop2 IHC in intestinal lesions from the genetic mouse model (Fig. 2D). While greater chromatin accessibility was found at fetal genes upon Apc inactivation (fig. S5C), new accessibility was observed at the *Tacstd2* genomic locus (Fig. 2G). These analyses suggest that expanded cell state plasticity by gaining access to and transcriptionally activating fetal intestinal genes is associated with intestinal neoplasia.

We observed that regenerative and fetal-intestinal gene signatures appeared to occupy relatively distinct compartments of the AbSC cluster, which was confirmed by a coexpression analysis of *Tacstd2* and *Ly6a* (fig. S5D), higher resolution re-clustering of AbSCs (fig. S5E), and relative nonoverlapping Sca-1 and Trop2 IHC in tdT+ intestinal lesions (Fig 2D). Pseudotime analysis of AbSCs indicated that regeneration transcriptional activity preceeded emergence of fetal intestinal gene expression (Fig. 2H). To examine this possibility, we performed scRNA-seq on tdT^+^ cells isolated from *Lgr5^Cre^;Apc^f/f^;R26^tdT^* mouse intestines 10 days following tamoxifen-induction (fig. S5F). As expected, AbSC activity was significantly elevated 10 and 28 days following Apc inactivation (fig. S5G). While *Ly6a* expression was stably elevated at day 10 and 28 (fig. S5H), fetal intestinal gene expression, including *Tacstd2*, gradually increased over time (Fig. 2I and fig. S5I), consistent with regeneration preceding developmental reprogramming. These data also suggest that nongenetic factors are responsible for fetal gene activation following regenerative transcriptional activity.

### Carcinogen-induced colon neoplasia demonstrates developmental reprogramming

To define features of AbSC in an independent model of colon neoplasia, we performed scRNA-seq of a neoplastic lesion and paired normal sample from a mouse exposed to AOM/DSS. UMAP representation of single-cell gene expression profiles of epithelial cells revealed known cell types in the normal colon and a distict trancriptional profile in the colonic lesion (Fig. 3A). AbSC gene signature derived from our genetic model demonstrated strongest expression in a neoplastic cluster furthest from normal tissue profiles and weaker expression in the remaining neoplastic cells and normal ISCs (Fig. 3B). Neoplastic cells demonstrated greater *Ly6a* expression by scRNA-seq (fig. 6A) and Sca-1 staining by IHC, the latter of which showed specific expression in lesions without expression in adjacent or distal normal tissue (Fig. 3C and fig. S6B). Notably, expression of the fetal intestinal gene signature broadly (Fig. 3D) and *Tacstd2* specifically (Fig. 3E and fig. S6C) was restricted to neoplastic cells. Consistent with these results, strong Trop2 expression was only found in premalignat and malignant lesions (Fig. 3C), corresponding to elevated Sox9 expression and attentuation of differentiation markers. These results demonstrate that a carcinogen-induced colon neoplasia model demonstrates developmental reprogramming in neoplastic cells, particularly marked by Trop2 positivity.

**Figure 3.**
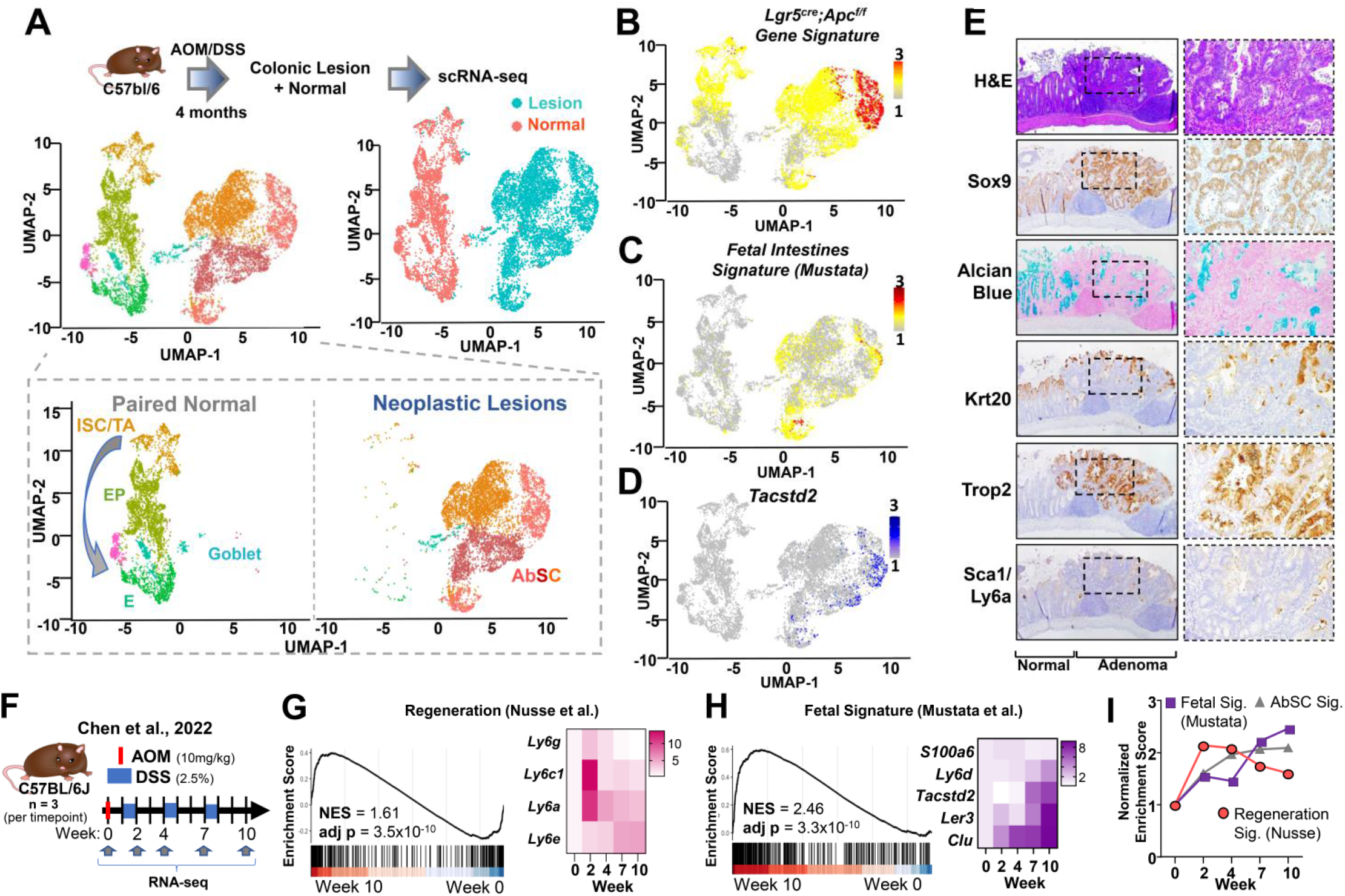
scRNA-seq of carcinogen-induced model of colonic neoplasia demonstrates developmental reprogramming. A. UMAP plot of epithelial cells from a colon lesion and adjacent normal epithelium of an AOM/DSS treated mouse model of colon carcinogenesis, colored by cell type (top left) and origin (top right). UMAPs from lesions and normal are separately depicted (bottom). ISC = stem cell, TA = transit amplifying, EP = enterocyte progenitor, E = mature enterocyte, G = goblet cell, AbSC = aberrant stem cell-like. B. Feature plot of Lgr5;Apc^KO^ gene signature on AOM/DSS UMAP. C. Feature plot of fetal-like gene signature (Mustata et al.) on AOM/DSS UMAP. D. Feature plot of *Tacstd2* (Trop2) expression on UMAP. E. Representative images of a high-grade dysplastic colonic lesion from an AOM/DSS treated mouse including hematoxylin and eosin (H&E) and Alcian blue staining as well as Sox9, Krt20, Sca-1 and Trop2 IHC. F. Experimental workflow for AOM/DSS induced colonic tumor model from Chen *et. al*., 2022 (GSE178145). Bulk RNA-seq of 3 biological replicates at indicated timepoints was analyzed. G. Gene set enrichment analysis (GSEA) of regeneration gene signature (Nusse et al., 2018) on AOM/DSS lesions at Week 10 vs controls (left) and heatmap of associated genes over time (right) (Chen *et.al*, 2022). H. GSEA of fetal-like intestinal gene signature (Mustata et al., 2013) (left) on AOM/DSS lesions at Week 10 vs controls (left) and heatmap of associated genes over time (right) (Chen *et.al*, 2022). I. Normalized enrichments scores for regeneration gene signature, fetal-like intestinal gene signature and AbSC gene signature over time in AOM/DSS data from (Chen *et.al*, 2022).

**Figure 4.**
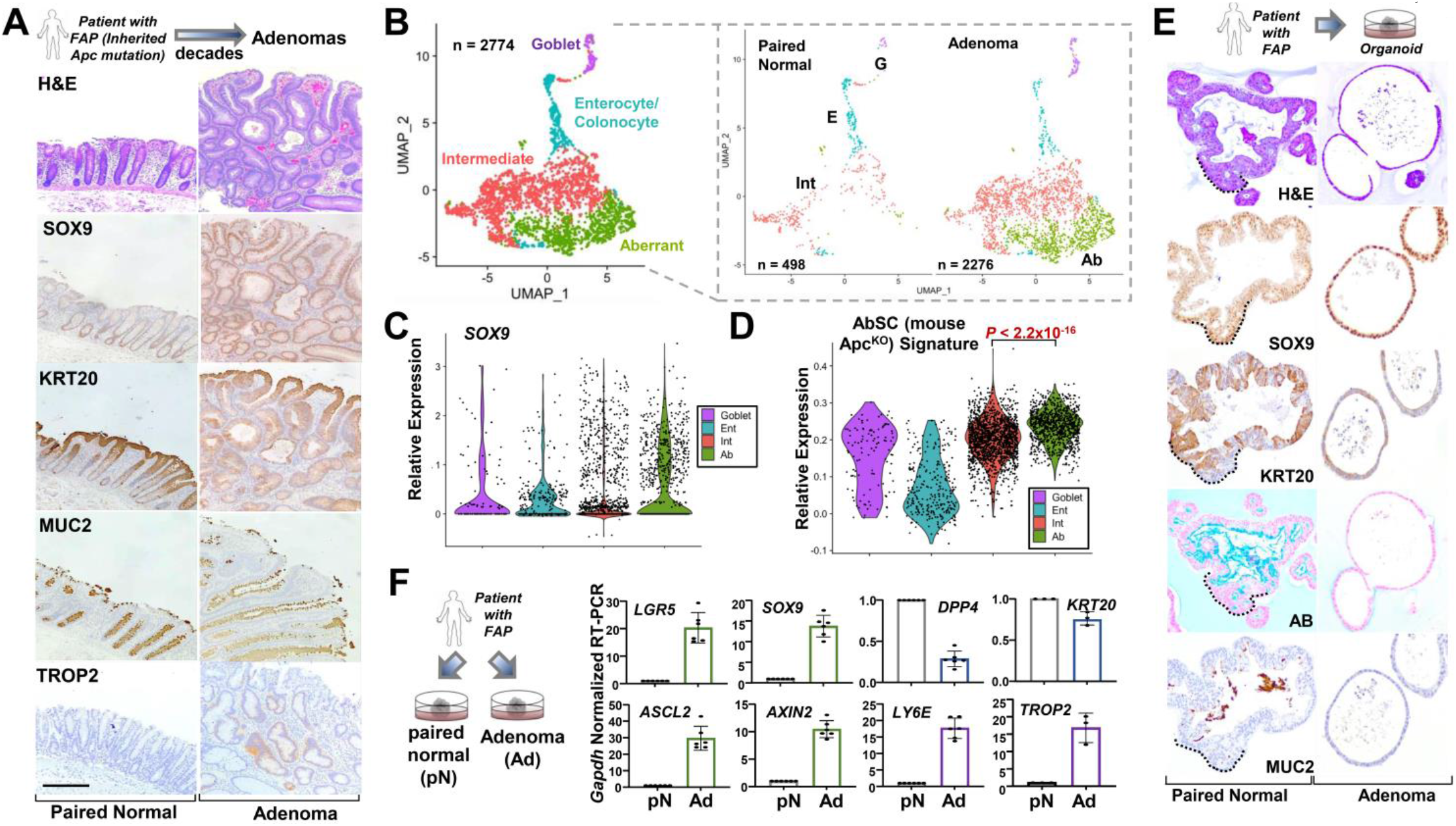
Characterization of human FAP adenomas by histopathology and scRNA-seq. A. Representative images of adenoma and normal adjacent colon tissue from patient with FAP patient including H&E staining SOX9, KRT20, and MUC2 IHC; scale bars = 250 μM. B. UMAP representation of scRNA-seq of human adenoma and paired normal indicating 4 distinct cell clusters; E = enterocyte, G = goblet, Int = intermediate, Ab = aberrant. C. Violin plot indicated *SOX9* expression in four different cell clusters D. Violin plot indicated expression of AbSC gene signature in four different cell clusters. *P*-value calculated by Wilcoxon rank sum test with Bonferroni correction. E. Representative images of organoids derived from adenoma and adjacent-normal tissue from patient with FAP including H&E and Alcian blue (AB) staining Sox9, Krt20, and Muc2 IHC; dotted line in paired normal sample indicates one of several crypts in the organoid F. mRNA expression of *SOX9*; stem cell marker *LGR5*; WNT pathway markers *AXIN2* and *ASCL2*; AbSC markers *LY6e, TROP2* and intestinal differentiation markers *KRT20, DPP4* in FAP organoids by qRT-PCR. Data expressed as mean ± S.D of three biological and two technical replicates.

**Figure 5.**
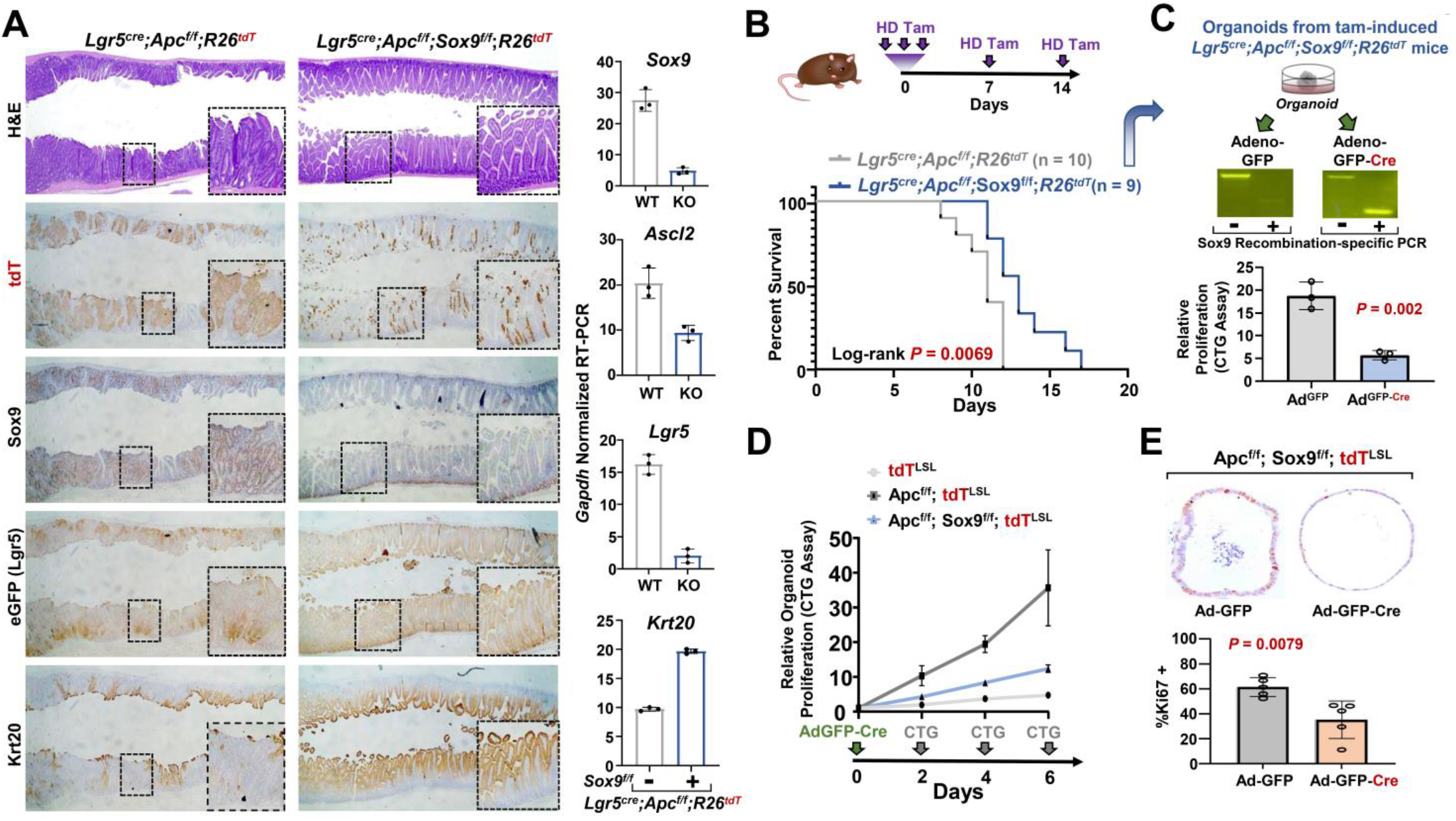
Sox9 is required for Apc^KO^ adenomas and organoids. A. Representative images of H&E staining and tdT, Sox9, Lgr5(eGFP) and Krt20 IHC from *Lgr5*-Apc^f/f^-tdT (control) and *Lgr5*-Apc^f/f^-Sox9^f/f^-tdT (experimental) mice small intestine; corresponding normalized mRNA expression of *Sox9, Ascl2, Lgr5*, and *Krt20*. Data expressed as mean ± S.D of three biological replicates B. Schematic of *in vivo* experimental design. Kaplan-Meier survival curve of *Lgr5*-Apc^f/f^-tdT (n = 10) and *Lgr5-* Apc^f/f^-Sox9^f/f^-tdT (n = 9) mice using high-dose (HD) tamoxifen (TAM) for induction and maintenance. Log-rank *P*-value = 0.0069. C. Organoids from *Lgr5*-Apc^f/f^-Sox9^f/f^-tdT mice were generated at experimental endpoint, treated with either Ad^GFP^ or Ad^GFP-Cre^ and subjected to *Sox9* recombination-specific PCR and a proliferation assay by CTG. Data expressed as mean ± S.D of three biological replicates. *P*-values calculated by two-sided Student’s t-test D. Proliferation of organoids derived from tdT, Apc^f/f^-tdT, and Apc^f/f^-Sox9^f/f^-tdT mice and infected with Ad^GFP-Cre^ at indicated time points by CTG. Data expressed as mean ± S.D of three biological replicates. E. Organoids derived from tdT, Apc^f/f^-tdT, and Apc^f/f^-Sox9^f/f^-tdT mice were infected with Ad^GFP^ or Ad^GFP-Cre^, formalin fixed, and then processed for histopathology. Representative images of Ki67 IHC. Quantification of %Ki67 positivity in 5-6 organoids per condition. *P*-values calculated by two-sided Student’s t-test

**Figure 6.**
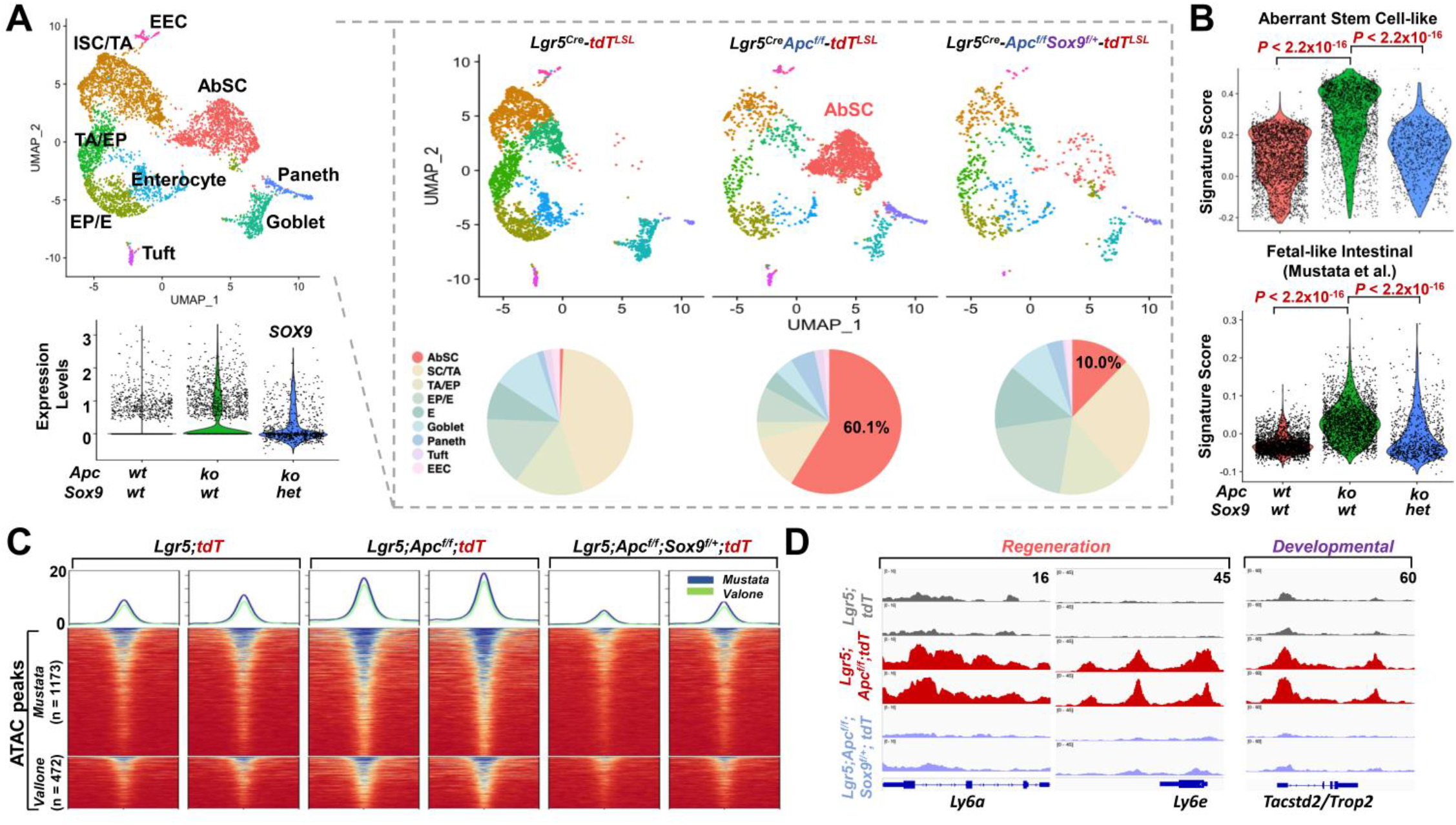
Sox9 suppression restricts AbSC and developmental reprogramming by scRNA-seq. A. UMAP representation of scRNA-seq data from tdT+ cells isolated by FACS from *Lgr5-tdT, Lgr5*-Apc^f/f^-tdT, and *Lgr5*-Apc^f/f^-Sox9^f/+^-tdT mice a month after tamoxifen induction colored by clusters and then separated by sample. Violin plot of normalized single cell Sox9 expression in each group. Pie charts indicating distribution of clusters in each sample. AbSC = aberrant stem cell-like, SC = stem cell, TA = transient amplifying, EP = enterocyte progenitor, E = enterocyte, EEC = enteroendocrine cell. B. Violin plot of normalized expression of AbSC gene signature (top) and fetal-like intestines gene signature (Mustata et al.) (bottom) in each group. *P*-value calculated by Wilcoxon rank sum test with Bonferroni correction. C. Heatmap depicting chromatin accessibility at fetal genes from 2 published signatures (Mustata et al. and Vallone et al.) in tdT+ cells flow-sorted from the intestines of *Lgr5-tdT* (n = 2), *Lgr5*-Apc^f/f^-tdT (n = 2), and *Lgr5*-Apc^f/f^-Sox9^f/+^-tdT (n = 2) mice. D. IGV screenshots depicting chromatin accessibility at *Ly6a, Ly6e* and *Tacstd2* genomic loci in tdT+ cells flow-sorted from the intestines of *Lgr5-tdT, Lgr5*-Apc^f/f^-tdT, and *Lgr5*-Apc^f/f^-Sox9^f/+^-tdT mice.

To validate and extend these findings, we analyzed a recently published bulk RNA-sequencing dataset derived from AOM/DSS treated mice at 4 different timepoints of colon neoplastic evolution (*49*) (Fig. 3F). Compared to untreated controls, transcriptional profiles of colonic lesions showed enhanced stem cell activity and impaired differentiation over time (fig. S6, D and E). Colonic lesions at week 10, the latest timepoint evaluated, showed significant enrichment of regeneration and fetal intestinal gene signatures compared to untreated controls (Fig. 3, G and H). The regeneration program peaked at week 2 and then gradually declined over time (Fig. 3, G and I). In contrast, the fetal intestinal gene signature gradually increased over time, showing strongest levels at week 10 (Fig. 3, H and I). These transcriptional program kinetics are consistent with histopathological evaluation of colonic lesions from our AOM/DSS model, which demonstrated robust Sca-1 expression in early compared to more advanced lesions, whereas Trop2+ expression followed the opposite pattern, with stronger staining in more advanced lesions compared to early ones (fig. S7). Furthermore, these results corroborate those found in our genetic model (Fig. 2, H and I), whereby developmental reprogramming lags behind regeneration activity in neoplastic progression (Fig. 3I).

### Human FAP adenoma and derivative organoids display impaired differentiation and AbSC activity

To evaluate transcriptional abnormalities in human CRC initiation, we evaluated adenoma and paired normal from a 39 year-old patient with familial adenomatous polyposis (FAP), a hereditary condition in which a mutant copy of *APC* is inherited, hundreds to thousands of intestinal adenomas develop, and prophylactic colectomy is required to prevent progression to cancer(*50*). In agreement with our mouse models, colon adenomas displayed elevated expression of SOX9 and TROP2 compared to adjacent normal tissue (Fig. 4A). Markers of differentiated absorptive and secretory colonic cells were partially suppressed as shown by KRT20 and MUC2 IHC, respectively (Fig. 4A). This observation could reflect limited WNT pathway restriction due to hypomorphic mutant APC activity(*51*) or admixture of normal colonic tissue.

We next performed scRNA-seq on cryopreserved adenoma and paired normal tissue. UMAP representation of single-cell gene expression profiles revealed 4 epithelial clusters derived from adenoma and normal tissue (Fig. 4B and fig. S8A); however, unlike our mouse models and consistent with human histopathology, the separation between adenomatous and normal tissue was less distinct. To define these clusters, we first utilized established gene signatures to identify normal enterocyte and goblet cell clusters (Fig. 4, B and C, and fig. S8 B to D). We then defined the cluster found only in the adenoma sample as ‘Aberrant’ (Fig. 4B). The remaining cluster was labelled ‘Intermediate’ since (1) it was found in both normal and adenoma tissue and (2) it did not engender a normal cell type gene expression profile(*38*) (Fig. 4B). Reassuringly, SOX9 expression was highest in the Aberrant cluster followed by the Intermediate cluster, whereas KRT20 expression was lowest in these two clusters (Fig. 4C and fig. S8C). In general, expression of ISC signatures was greatest in the Aberrant cluster relative to Intermediate, enterocyte, and goblet clusters (figs. S9A and S8E), further building confidence in our classification. Notably, AbSC gene expression signature was greatest in the Aberrant cluster and adenoma tissue (Fig. 4D and fig. S8F). A subset of regeneration and fetal-like intestinal signatures and genes, including *TACSTD2*, were elevated in the Aberrant cluster (figs. S9B, and S8, G and H). We validated these findings using published scRNA-seq data from patients with CRC (n = 9) using normal colon (n=3) as a comparator (fig. S10, A to E). Together, these data indicate that human adenomas demonstrate a partial block in differentiation and activation of an AbSC program involving developmental programs.

We next evaluated organoids derived from adenoma and normal colonic tissue from the same patient with FAP, asking whether features of AbSC activity are captured in this three-dimensional culture system that requires supplemented WNT3A, R-spondin, and Noggin (WRN) conditioned media rich in stem cell promoting factors(*52*). Normal colonic cultures contained organoids that appeared folded by phase contrast and generated crypts with high SOX9 expression, absent KRT20 expression, and absent MUC2 expression/Alcian blue (AB) staining (Fig. 4E, dotted line). These organoids also displayed differentiated regions with low or no SOX9 expression, high KRT20 expression, intermittent MUC2 expression, and strong AB staining (Fig. 4E). In contrast, most organoids in adenomas cultures (>95%) did not display differentiated regions as indicated by unfolded, spheroid morphology by phase contrast and weak KRT20 levels and absent MUC2 expression by IHC. Rather, adenoma organoids expressed uniform high SOX9 levels (Fig. 4E). These findings were confirmed in a broader survey of stem cell/WNT (*LGR5, SOX9, ASCL2, AXIN2*) and regeneration/developmental genes (*LY6E, TACSTD2*), which demonstrated markedly higher mRNA expression in adenoma compared to normal organoid cultures (Fig. 4F). *Ly6a* was not profiled as it does not have a direct human homolog. mRNA expression of colonic differentiation markers *DPP4* and *KRT20* were correspondingly reduced, albeit modestly due to enhanced stem cell cues conferred by WRN media (Fig. 4F).

We validated these results in Apc^WT^ and Apc^KO^ organoids derived from our genetic mouse model (fig. S9, C to E). After confirming Apc inactivation in tdT^+^ organoids conferred niche independence (the ability to grow without WRN media) (fig. S9D), we performed bulk RNA-sequencing. In addition to activation of stem cell genes and partial reduction in intestinal differentiation, Apc^KO^ organoids demonstrated robust activation of developmental intestinal genes, especially *Tacstd2* (fig. S9E). Collectively, these data suggest that human adenoma and mouse Apc^KO^ organoids display features of developmental reprogramming as part of an AbSC transcriptional program.

### Sox9 is required for intestinal neoplasia initiation

Based on its (1) enriched transcriptional activity in AbSCs, (2) stronger expression in fetal intestines compared to adult intestines, and (3) ability to functionally block differentiation in models of CRC(*9*), we hypothesized that Sox9 is required for CRC initiation. To evaluate this hypothesis, we bred *Lgr5^Cre^;Apc^f/f^;R26^tdT^* mice with *Sox9^f/f^* mice(*53*) to homozygosity, enabling conditional biallelic deletion of Sox9 in the Apc^KO^ setting upon tamoxifen induction. Experiments were initiated with three daily tamoxifen injections followed by weekly single injection maintenance treatments, attempting to force robust genetic recombination. Successful Sox9 inactivation prevented tdT^+^ adenoma formation, reduced mRNA levels of stem cell genes (*Ascl2, Lgr5*) and maintained intestinal homeostasis with crypt restricted Lgr5^eGFP+^ stem cells and Krt20^+^ differentiated villi (Fig. 5A). While *Lgr5^Cre^;Apc^f/f^;Sox9^f/f^;R26^tdT^* mice survived significantly longer than *Lgr5^Cre^;Apc^f/f^;R26^tdT^* controls (Fig. 5B), they eventually succumbed to adenoma burden. These adenomas invariably expressed Sox9 despite tdT^+^ activation and Apc deletion, suggesting escape of Sox9 deletion, which we confirmed by PCR of flow-sorted tdT^+^ cells demonstrating a significant fraction of unrecombined Sox9 by (fig. S11A). To show that tdT^+^ adenomas from *Lgr5^Cre^;Apc^f/f^;Sox9^f/f^;R26^tdT^* mice were still dependent on Sox9, we generated organoids at the experimental endpoint and subjected them to *ex vivo* recombination using specific adenovirus (Ad) infections. Compared to organoids treated with mock Ad^GFP^ controls, Ad^GFP-Cre^ treated organoids displayed reduced proliferation by CellTiterGlo© (CTG), which corresponded to robust Sox9 inactivation by recombination (Fig. 5C). We confirmed these results by performing a 10-day short-term experiment, isolating organoids from three mice from each genotype, and subjecting them to Ad^GFP-Cre^ infections followed by CTG (fig. S11B). Collectively, these experiments indicate that successful Sox9 inactivation prevents intestinal Apc^KO^ adenomas using genetic engineered mice and derivative organoids.

We next established colon organoids without endogenous Cre recombinase for controlled *ex vivo* Ad^GFP-Cre^ experiments, seeking further validation of Sox9 dependency in colonic adenomas. Colon organoids from *R26^tdT^,Apc^f/f^;R26^tdT^*, and *Apc^f/f^;Sox9^f/f^;R26^tdT^* mice were infected with Ad^GFP-Cre^ and subjected to proliferation assays by CTG. While Apc inactivation led to a 7.5-fold increase in organoid proliferation, concomitant Sox9 deletion reduced this proliferative advantage by 3-fold (Fig. 5D). To support these findings, we performed tdT^+^ quantification by FACS as a surrogate for cell viability following Ad^GFP-Cre^ treatment. tdT was activated in 54% of *Apc^f/f^;R26^tdT^* organoids 4 days after Ad^GFP-Cre^ infection, which was reduced by 4-fold to 13% in *Apc^f/f^;Sox9^f/f^;R26^tdT^* organoids (fig. S11C); similar results were achieved with lentiviral Cre activation (fig. S11D). Finally, compared to Ad^GFP^ treated controls, Ad^GFP-Cre^ treated *Apc^f/f^;Sox9^f/f^;R26^tdT^* organoids demonstrated impaired proliferation by Ki67 IHC, which was not observed in *R26^tdT^* or *Apc^f/f^;R26^tdT^* organoids (Fig. 5E and fig. S11E). *Apc^f/f^;Sox9^f/f^;R26^tdT^* organoids with more efficient Ad^GFP-Cre^ infection marked by increased tdT activation demonstrated reduced Sox9 expression and lower proportion of Ki67^+^ cells relative to poorly infected ones (fig. S11E, red vs. gray arrow). These experiments demonstrate a requirement for Sox9 in adenoma formation using a genetically engineered mice and organoids.

### scRNA-seq reveals Sox9 is required for AbSC

To determine whether Sox9 is required for AbSC transcriptional activity, we performed scRNA-seq on FACS-isolated tdT+ cells from *Lgr5^Cre^;Apc^f/f^;Sox9^f/+^;R26^tdT^* mice one month after tamoxifen induction. We chose *Lgr5^Cre^;Apc^f/f^;Sox9^f/+^;R26^tdT^* mice to analyze tdT^+^ cells with consistent and reliable reduction in Sox9 expression given our difficulty to isolate viable tdT^+^ cells with efficient homozygous deletion of Sox9 (Fig. 5, B and C). Reassuringly, viable tdT^+^ cells from *Lgr5^Cre^;Apc^f/f^;Sox9^f/+^;R26^tdT^* mice demonstrated a considerable reduction in Sox9 levels compared to *Lgr5^Cre^;Apc^f/f^;R26^tdT^* mice, which displayed higher Sox9 expression relative to *Lgr5^Cre^; R26^tdT^* controls (Fig. 6A and fig. S12, A and B). UMAP representation of single-cell gene expression profiles revealed clusters representing the normal epithelial cell types and AbSCs (Fig. 6A and fig. S12C). Notably, single copy deletion of Sox9 reduced the percentage of AbSCs from 60% to 10% in tdT^+^ Lgr5-Apc^KO^ cells (Fig. 6A and table S3) and significantly reduced AbSC gene signature activity (Fig. 6B), indicating a requirement for Sox9 in maintaining an AbSC transcriptional program. ISC gene signatures were also reduced in tdT^+^ cells from *Lgr5^Cre^;Apc^f/f^;Sox9^f/+^;R26^tdT^* mice (Fig. 6B, and fig. S12, D and E). The 6-fold reduction in AbSC cells was accompanied by restoration of a normal distribution of ISC and differentiated intestinal cell types (Fig. 6A and table S3). We also observed reduced expression of IFN-γ, regeneration signatures, including *Ly6a*, and fetal-like intestinal gene signatures, including *Tacstd2*, in tdT^+^ cells from *Lgr5^Cre^;Apc^f/f^;Sox9^f/+^;R26^tdT^* mice (fig. S11, F to K). These results were confirmed *R26^tdT^,Apc^f/f^;R26^tdT^*, and *Apc^f/f^;Sox9^f/f^;R26^tdT^* organoids using *ex vivo* Ad^GFP-Cre^ infections followed by RT-PCR (fig. S13). ATAC-seq profiling revealed that Sox9 inactivation led to reduced chromatin accessibility at regeneration and fetal genes in Apc^KO^ cells (Fig. 6C and fig. S14A), specifically at the *Ly6a, Ly6e* and *Tacstd2* genomic loci (Fig. 6D and fig. S14B), indicating rescue of expanded cell state plasticity. Collectively, these results demonstrate a requirement of Sox9 in AbSC activity, aberrant cell state plasticity, and fetal intestinal gene reactivation.

### SOX9 knockdown in FAP adenoma organoid induces differentiation

To evaluate SOX9 activity and necessity in human colonic neoplasia, we utilized our FAP adenoma organoid model (Fig. 4E). We first defined genomic SOX9 binding pattern in paired normal colon and adenoma organoids by CUT&RUN. A global increase in SOX9 binding was observed in adenoma organoids compared normal colon organoids (fig. S15A). Greater SOX9 binding was observed at ISC and AbSC genes in adenoma compared to normal colon organoids (Fig. 7A and fig. S15B); while many sites are shared, there are greater than 4-fold unique high-confidence SOX9 binding at AbSC genes in adenoma compared to normal (Fig. 7B). Consistent with selective ISC activity upon Apc inactivation, we noticed that SOX9 binding at ISC gene *LRIG1* was unchanged in adenoma compared to normal colon organoids (fig. S15C), whereas greater SOX9 binding was found at a *PROM1* intronic enhancer (fig. S15C), validating our previous work (*9*) and supporting SOX9-mediated AbSC activity in neoplasia. We also found greater SOX9 binding at both Mustata and Vallone fetal genes as well as Nusse regenerative genes in adenoma compared to normal colon organoids (Figure 7, B and C, and fig. S15D-F). Consistently, stronger SOX9 binding was found at fetal genes, including *TACSTD2* and *CLU*, in adenoma compared to normal colon organoids (Figure 7D). Despite the enrichment in adenoma organoids, we think there is considerably more SOX9 binding at fetal genes in normal colon organoids than expected, likely due to strong and sustained influences of WNT/R-Spondin/Noggin conditioned media.

**Figure 7.**
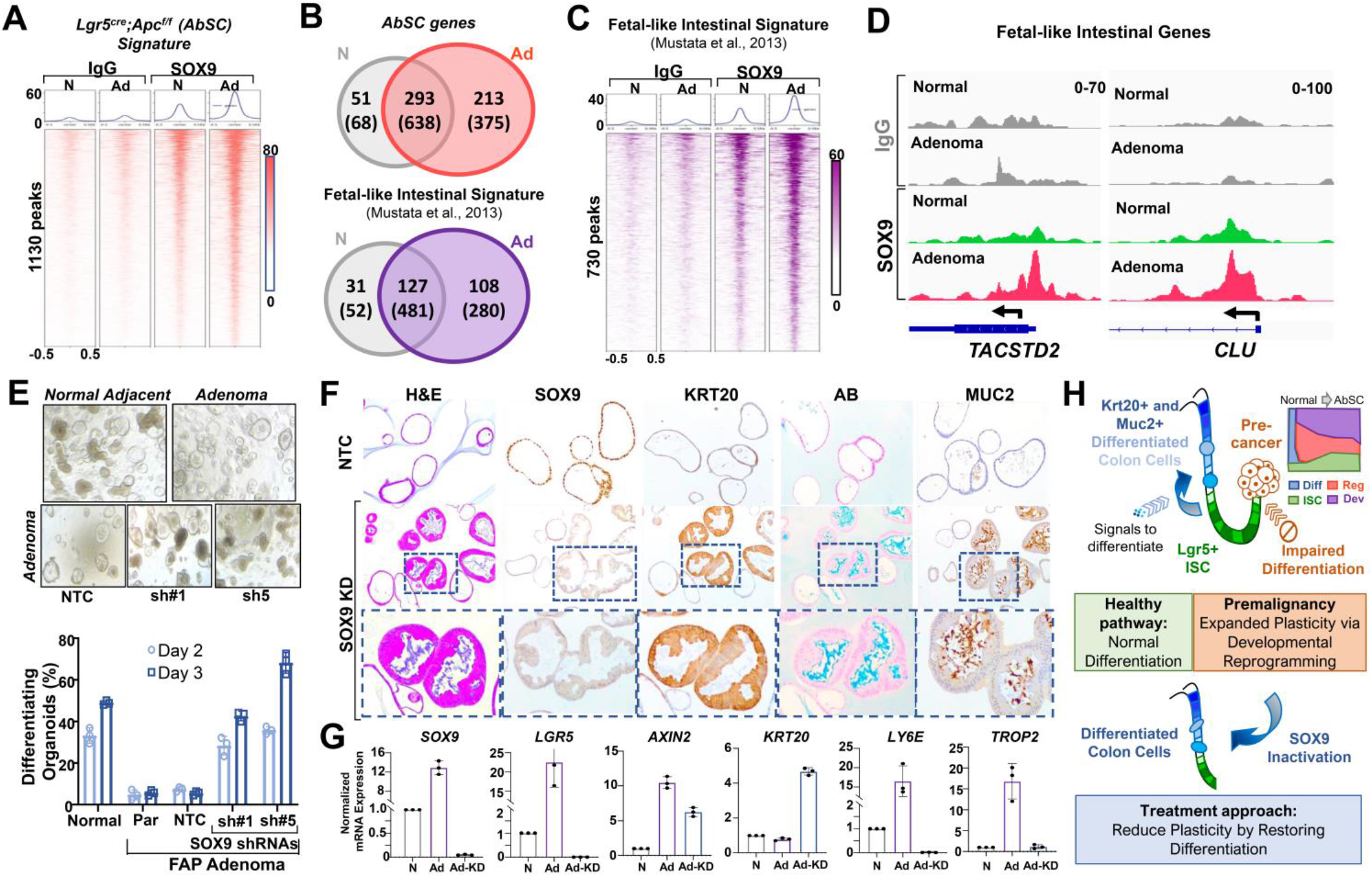
SOX9 KD impairs fetal reprogramming and induces differentiation in FAP adenoma organoids. A. Heatmap of SOX9 binding associated with AbSC genes (1130 peaks) in normal colon (n = 4) and adenoma (n = 4) organoids from a patient with FAP by CUT&RUN. B. Venn diagram of high-confidence SOX9 bound genes (peaks) using LFC ≥ 0.5 compared to IgG control in normal colon and adenoma organoids for genes in AbSC (top) and Mustata fetal-like intestines (bottom) C. Heatmap of SOX9 binding associated with Mustata fetal-like intestinal (730 peaks) in normal colon and adenoma organoids by CUT&RUN. D. SOX9 binding at fetal intestinal genes *TACSTD2* and *CLU* in normal colon and adenoma organoids by CUT&RUN. E. Phase contrast images depicting differentiation phenotype (folded) of organoids derived from normal colon and adenoma organoid cultures (top row); adenoma organoids expressing non-targeting control (NTC), or two distinct shRNAs against SOX9 (bottom row). Quantification of differentiating organoids in indicated cultures at day 2 and 3; Par = parental. F. Representative images of H&E, Alcian blue (AB) staining and SOX9, KRT20, and MUC2 IHC of NTC and SOX KD FAP adenoma organoids. G. Normalized mRNA expression of *SOX9, LGR5, AXIN2, KRT20, LY6E, TROP2* in indicated organoids: normal adjacent mucosa (N), adenoma-NTC (Ad) and adenoma-SOX9 KD (Ad-KD). H. Schematic summarizing AbSC transcriptional program and developmental reprogramming obstructing intestinal differentiation in CRC initiation and the ability of SOX9 suppression to reverse these effects.

We next asked whether SOX9 is required for AbSC activity in human FAP adenoma organoids. Human adenoma organoid cultures showed few differentiating organoids (~5%), which appear as folded structures by phase contrast, whereas organoid cultures derived from normal adjacent tissue displayed evidence of a differentiating population that amounted to about 50% by day 3 (Fig. 7E and fig. S16A). Using this system, we examined whether shRNA mediated SOX9 knockdown (KD) can force differentiation in human adenoma organoids. Stable SOX9 KD shifted the adenoma culture to consist of 40-70% differentiating organoids, representing up to a 14-fold increase compared to parental and non-targeting control (NTC) by day 3 (Fig. 7E, and fig. S16, B and C). Histopathology of fixed organoids demonstrated that SOX9 KD induced robust KRT20 expression by IHC and mucin production by AB staining and MUC2 IHC (Fig. 7F). Consistently, SOX9 KD reduced stem cell activity (*LGR5, SOX9, AXIN2*) and regenerative/developmental reprogramming (*LY6E, TROP2*) while inducing differentiation (*KRT20*) (Fig. 7G). In contrast, SOX9 KD led to a modest, statistically insignificant increase in differentiation of normal colon organoids, which corresponded to induction of KRT20 without impact on fetal genes (fig. S16, D and E). These data indicate that human FAP adenomas depend on SOX9 to maintain developmental reprogramming and prevent differentiation (Fig. 7H).

## DISCUSSION

Our study indicates that reactivation of a fetal intestinal program is a critical component of an aberrant transcriptional state that restricts differentiation and expands cell state plasticity during CRC initiation. *APC* is the most frequently mutated gene in CRC(*19*). In the intestines, *Apc* inactivation leads to unrestricted WNT activity and ISC expansion, which contributes to adenoma formation. However, whether WNT activation by APC loss-of-function leads to normal ISC transcriptional activity could not be resolved using previous methods. By applying scRNA-seq to an Apc^KO^ genetic mouse model, we found that Apc inactivation leads to a transcriptional cell state that is distinct from normal ISCs; we therefore refer to this program as an aberrant rather than a hyperactive stem cell-like state. While upregulation of the WNT pathway and ISC activity is part of the AbSC program, these alone do not capture the complete cadre of deviant consequences following Apc inactivation. A unique transcriptional program that consists of inappropriate interferon related signaling, sustained regeneration, and re-engagement of developing fetal intestinal genes was found in a subset of Apc^KO^ intestinal epithelial cells. Interestingly, there appears to be convergence of two of these transcriptional programs on *Ly6a*, which has been shown to be a target of IFN signalling in pathological states in the intestines such as colitis(*54*) and infection(*42*) as well as implicated in epithelial regeneration following injury(*55*) or stem cell ablation(*42*, *56*). Regenerative transcriptional activity is transient during pathological states, such as infection, and expresssion of fetal genes is temporally restricted during development. However, expression of these programs following Apc inactivation is sustained and may have consequences that permit neoplastic progression. In fact, it appears that persistant regeneration activity precedes and may enable emergence of developmental gene expression. In agreement with these results, open chromatin was found at regenerative genes in line with the potential need to express these programs with prompt kinetics, whereas new accessibility was associated with fetal genes, potentially explaining its delayed expression following Apc inactivation.

Reactivation of genes that operate during fetal development has been observed in CRC(*57*) and other cancers(*58*, *59*); however, our data indicate that reactivation of developmental genes can be traced to premalignant disease prior to malignant transformation. Identification of fetal genes that are consistently activated in premalignant colonic lesions therefore carry significant diagnostic value as their expression would normally be silenced in adult tissue under homeostatic conditions; the difficulty may arise in the specifity of such a diagnostic marker if transiently expressed during infectious, inflammatory, or other non-neoplastic pathologic states. Carcinoembryonic antigen, better known as CEA, is a CRC biomarker that has proven useful in the perioperative and disease recurrence settings(*60*, *61*). Accessing silenced regions of the genome that were once active during pluripotent development would intuitively serve the faculty of expanding cell state plasticity and adaptive fitness among other advantageous properties for neoplastic growth(*62*) and metastasis(*63*). Importantly, developmental reprogramming is a nongenetic feature that would confer clonal fitness through epigenetic inheritance. Defining the precise functional importance of aberrant cell state plasticity in cancer(*6*) and fetal intestinal program reactivation in CRC initiation(*57*) will require deeper characterization.

One unifying consequence of the aberrant transcriptional state involves impaired intestinal differentiation, although our experiments do not necessarily distinguish between blocked differentiation and de-differentiation, two often indistinguishable mechanisms leading to expanded cell state plasticity(*6*). Since the intestine is the most rapidly renewing epithelium in the human body, any genomic alteration or transcriptional state that promotes neoplasia must counteract the natural tissue turnover. A mutation that does not promote intestinal tissue fixation is unlikely to have neoplastic consequences, at least during cancer initiation. Indeed, beyond genetic mouse models that delete or suppress *Apc*, random mutagenesis leads to premalignant and malignant lesions with a similar molecular pattern of impaired differentiation. Adenomas from a patient with an inherited *APC* mutation demonstrated attenuation of intestinal differentiation rather than a complete block. There is evidence that adenomas in patients with FAP select for *APC* mutations that retain partial function, which is postulated to confer optimal β-catenin activity to facilitate cancer initiation(*64*). In agreement with these genomic findings, biochemical reconstruction of the destruction complex demonstrated that truncated APC mutants are functional hypomorphs that retain limited WNT restricting capacity(*51*), likely explaining their ability to partially obstruct differentiation, impart clonal fitness, and yield insidious neoplastic benefits.

Given that preventing differentiation is a hallmark of CRC initiation, the translational question is whether restoring or inducing differentiation can serve as a therapeutic strategy, one that has proven successful for a subset of leukemias(*6*, *65*). The proof of concept of this strategy for CRC lies in studies that reintroduced functional APC and demonstrated restoration of regulated WNT signaling, induction of intestinal differentiation, and inhibition of cancer growth(*27*–*29*). Furthermore, disrupting WNT-dependent stem cell programs by eliminating Lgr5+ stem cells (*30*, *31*) or ectopically expressing pro-differentiation transcription factors such as HOXA5(*66*) impaired tumor growth and metastasis in preclinical models. Translating these important studies into effective therapeutics remains the challenge. One possibility is that interfering with normal ISC activity is insufficient given the additional aberrant faculties endowed by Apc inactivation. SOX9 is not only important for adult crypt biology and ISC activity but also appears to be a critical driver of the AbSC state. Sox9 was previously shown to be required for intestinal regeneration, a component of AbSC transcriptional program, following radiation injury (*67*). Compromised Apc function led to elevated Sox9 expression, which may approach levels of Sox9 found in fetal intestinal development (*1*) and enable reactivation of developmental genes through pioneering activity (*68*), especially in the setting of greater chromatin accessibility. SOX9 may therefore serve as a better target than other ISC drivers as disrupting its activity will also interfere with aberrant developmental reprogramming. Indeed, genetic inactivation of *Sox9* in our mouse model and suppression in human FAP adenomas not only reduces ISC gene signatures but also fetal intestinal transcriptional programs. Beyond preventing adenoma formation, SOX9 suppression also disrupts growth of established neoplastic murine organoids and human cancers by inducing intestinal differentiation(*9*). As such, SOX9 is required for cancer initiation and maintenance; it can therefore be targeted as a preventative strategy in hereditary polyposis syndromes and as a treatment approach in sporadic CRC.

## Supplementary figures 1-16

**fig. S1.**
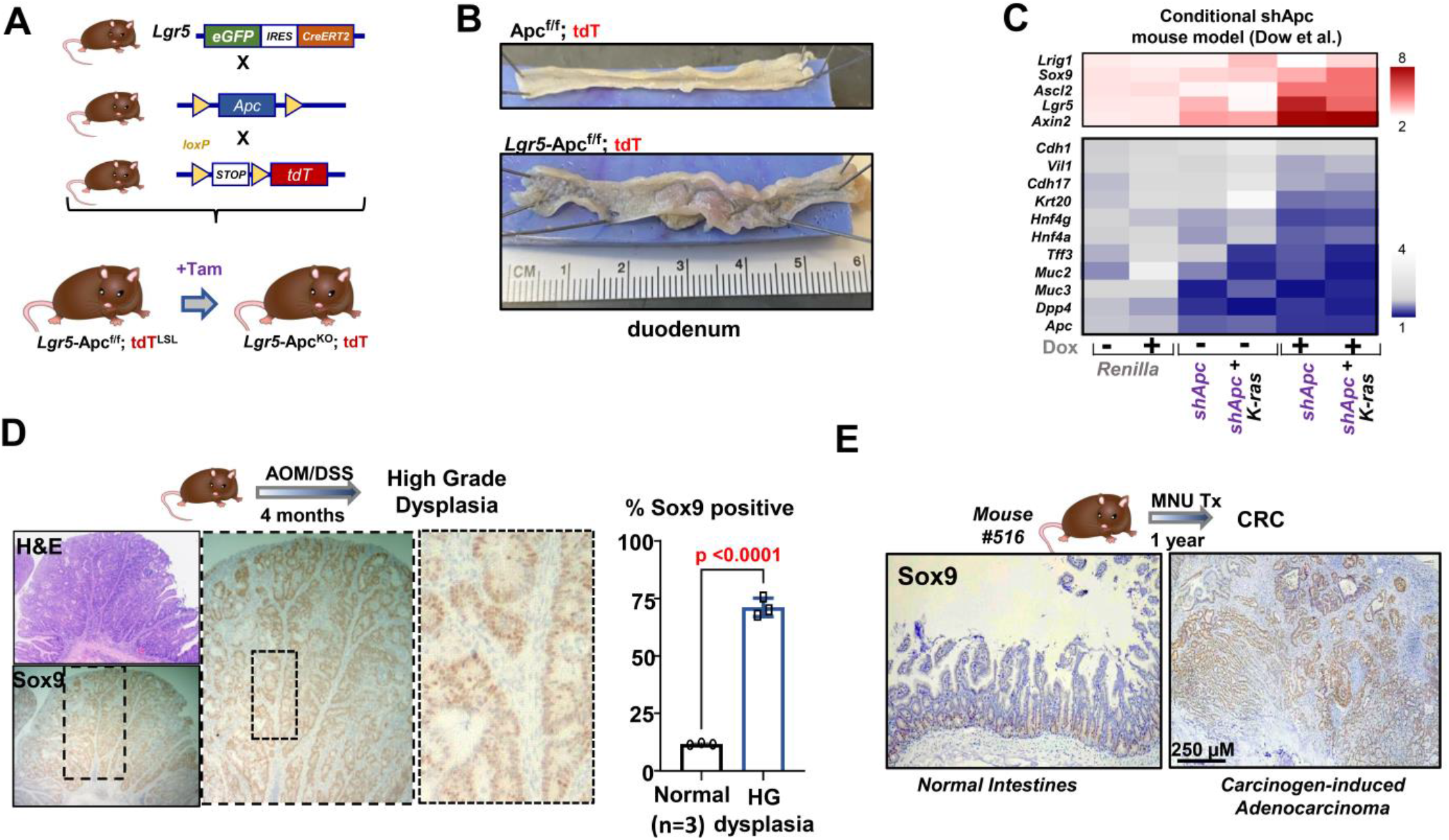
Elevated expression of Sox9 and reduced expression of differentiation genes in murine models of intestinal neoplasia. A. Diagram showing breeding scheme to achieve *Lgr5*-Apc^f/f^-tdT mice for conditional *Apc* inactivation. B. Representative images showing gross anatomy of duodenal intestines from indicated mice. C. Heat map of intestinal stem cell (red) and differentiation (blue) genes from previously published mouse models (Dow et al., 2015). RNA-seq data from the following three mice with and without dox are presented: shRNA against Renilla, Apc (shApc) alone or in combination with mutant K-ras allele. D. High magnification representative images of Sox9 overexpression in dysplastic crypts of AOM/DSS model (left); quantification of Sox9 positivity from three lesions. E. High magnification representative images of Sox9 overexpression in carcinoma from MNU model; scale bar = 250 μM.

**fig. S2.**
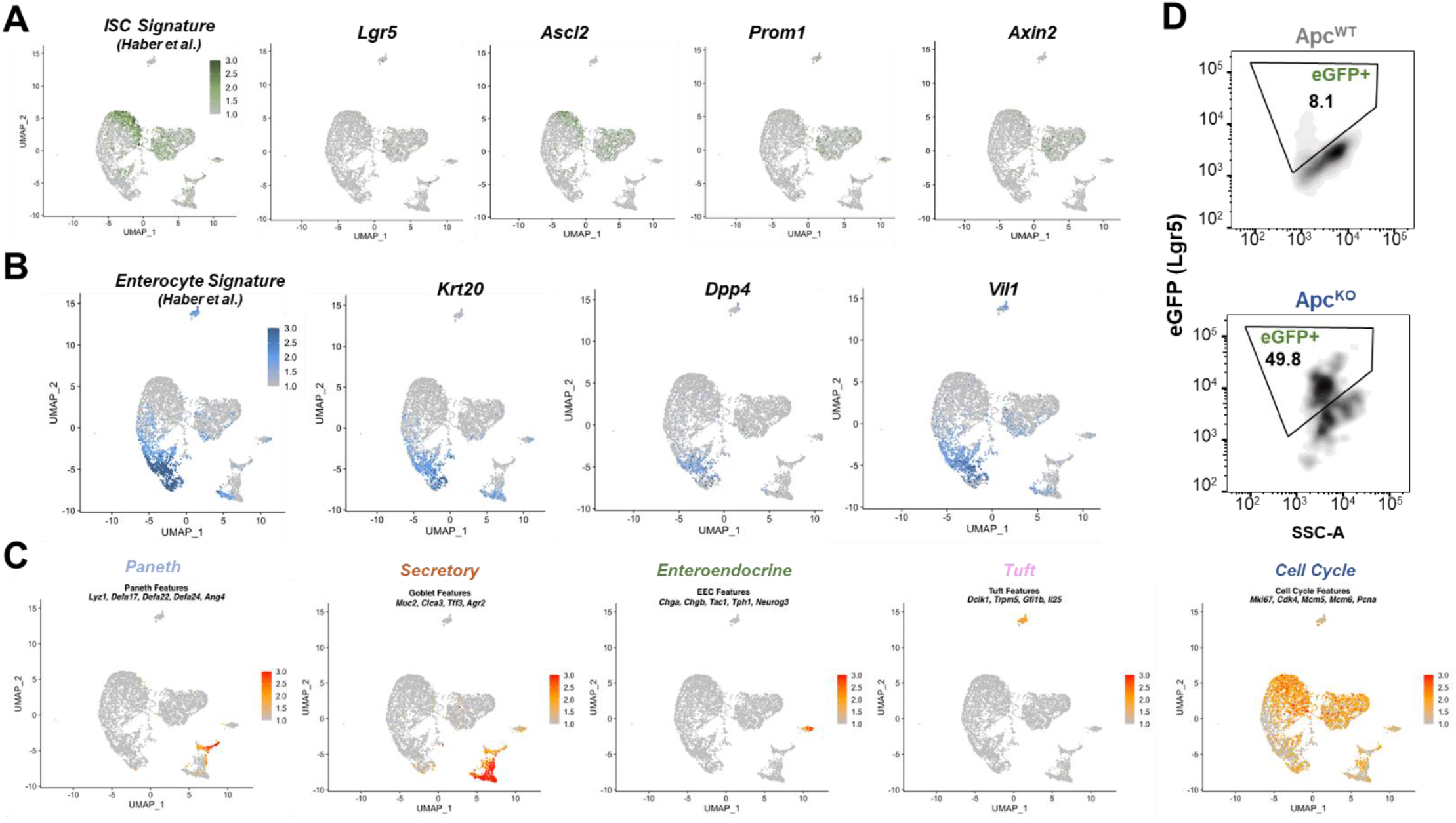
Single cell transcriptional profiling of tdT+ cells from Apc^KO^ genetic mouse model. A. UMAP representation of experiment in Figure 2A colored by ISC signature (Haber et al.), individual stem cell genes (*Lgr5, Prom1, Olfm4, Ascl2*), Wnt pathway gene *Axin2*. B. UMAP representation of experiment in Figure 2A colored by Enterocyte signature (Haber et al.), individual enterocyte genes (*Krt20, Dpp4, Vil1*). C. UMAP representation of experiment in Figure 2A colored by Paneth cell gene signature, Secretory cell gene signature, enteroendocrine gene signature, tuft cell gene signature, and cell cycle gene signature. D. Distribution of eGFP^+^ cells (Lgr5 expression) from tdT+ cells isolated by FACS from Lgr5;Apc^f/+^;tdT (control) and Lgr5;Apc^f/f^;tdT (experimental) mice.

**fig. S3.**
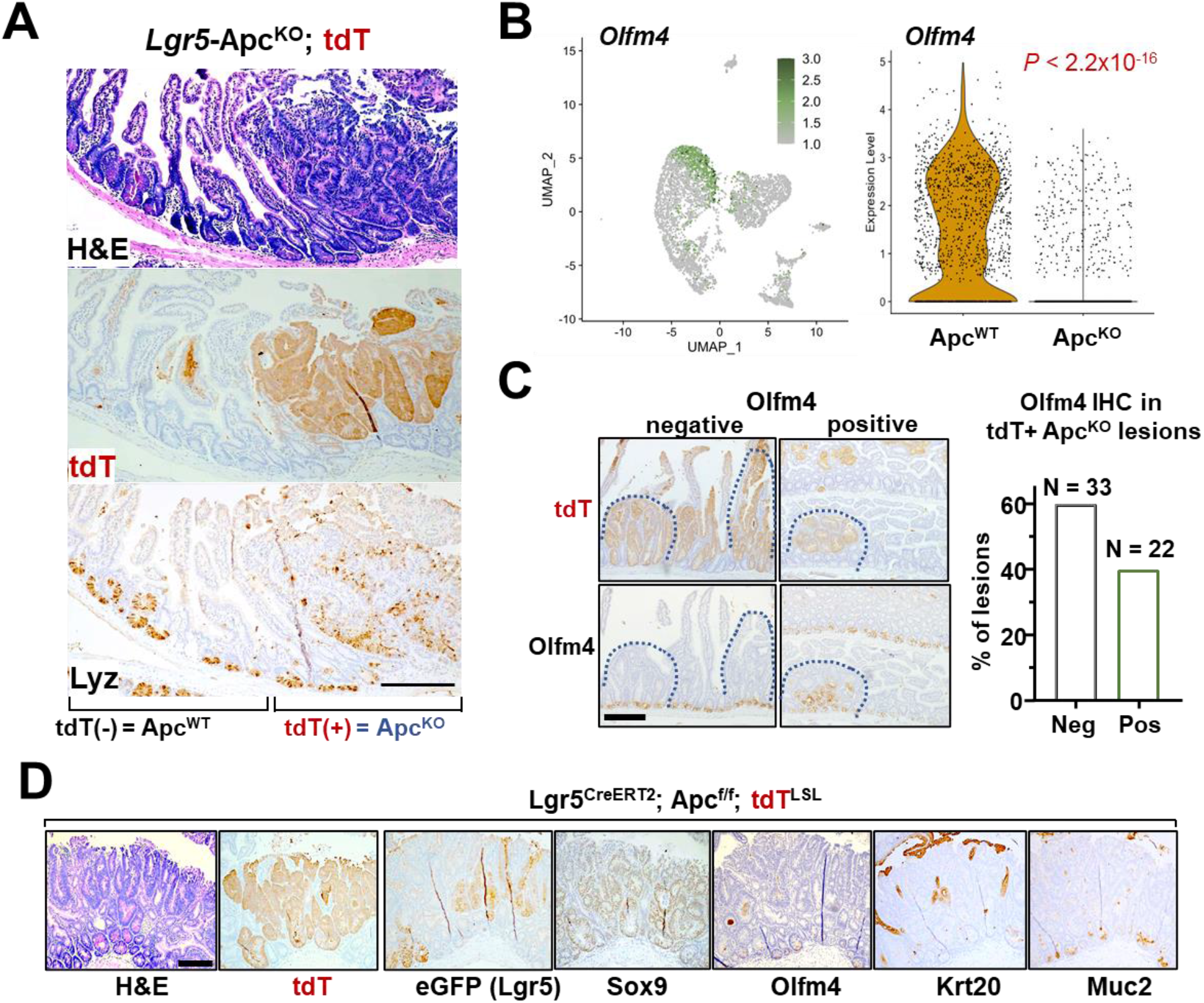
Apc^KO^ lesions demonstrate ectopic Paneth cell and selective ISC. (A) Representative images of intestinal lesions from *Lgr5*-Apc^f/f^-tdT mice including H&E staining, tdT and Lysozyme (Lyz) IHC. Adjacent normal (Apc^WT^) and adenoma (Apc^KO^) regions as marked by tdT and labeled; scale bars = 250 μM. (B) UMAP representation colored by *Olfm4* expression (left); violin plot of *Olfm4* expression in indicated samples (right). *P*-value calculated by Wilcoxon rank sum test with Bonferroni correction. (C) Representative images of intestinal lesions from *Lgr5*-Apc^f/f^-tdT showing tdT and Olfm4 IHC. Quantification of lesions that are positive (n = 33) and negative (n = 22) for Olfm4 expressed as a percentage of total tdT lesions; scale bars = 250 Mm (D) Representative images of intestinal lesions from *Lgr5*-Apc^f/f^-tdT showing H&E, tdT, eGFP (Lgr5), Sox9, Olfm4, Krt20, Muc2 IHC.

**fig. S4.**
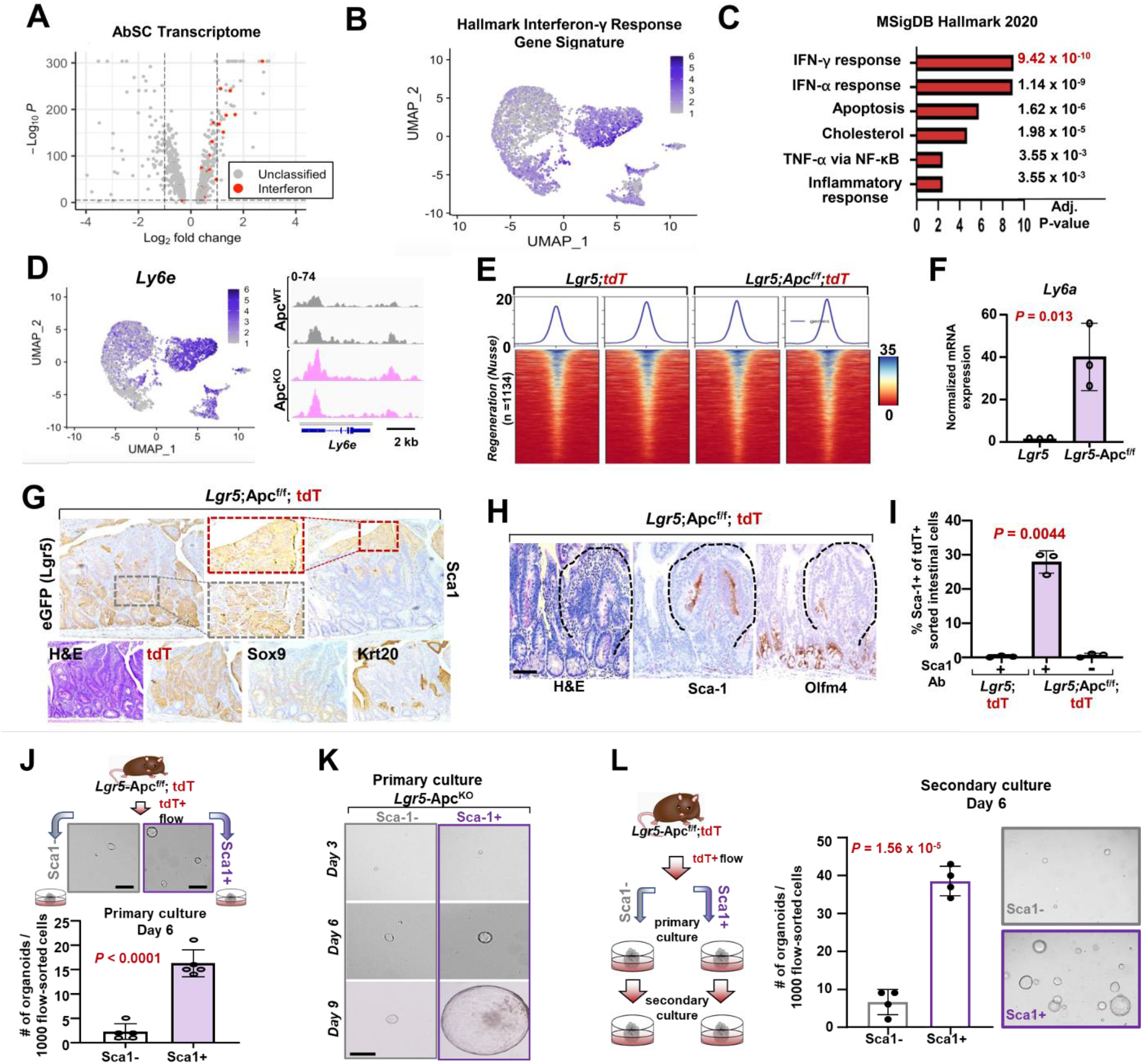
IFN signaling and regeneration genes are upregulated in AbSC program. A. Volcano plot representing differentially expressed genes from AbSC cluster. Genes associated with interferon signaling (red) are highlighted. B. Normalized expression of IFN-γ response gene signature on UMAP plot. C. MSigDB Hallmark significantly enriched gene-sets in AbSC cluster (EnichR) D. Normalized expression of *Ly6e* on UMAP plot (left) and chromatin accessibility at *Ly6e* genomic locus in tdT+ cells isolated by FACS from *Lgr5-tdT* and *Lgr5*-Apc^f/f^-tdT mice by ATAC sequencing (right). E. Heatmap depicting opening of regeneration genes in Lgr5-Apcf/f-tdT mice compared to Lgr5-tdT mice intestines published signature (Nusse et al.). F. mRNA expression of Ly6a in tdT+ cells isolated by FACS from Lgr5-tdT and Lgr5-Apcf/f-tdT mice by qRT-PCR. G. Representative images of H&E staining and eGFP(Lgr5), Sca-1, Sox9, Krt20, and tdT IHC from *Lgr5*-Apc^f/f^- tdT mice; scale bars = 100 μM and 20 μM. H. Representative images of H&E staining and Sca-1 and Olfm4 IHC from *Lgr5*-Apc^f/f^-tdT mice; scale bars = 100 μM and 20 μM. I. Percentage of tdT+ cells that are Sca-1+ by FACS in Lgr5-tdT and Lgr5-Apcf/f-tdT mice in indicated conditions. J. Representative image (top) and quantification (bottom) of organoids formed on Day 6 after plating 1000 FACS-isolated tdT+/Sca-1+ and tdT+/Sca-1- cells from *Lgr5*-Apc^**f/f**^-tdT mice; mean ± S.D of three biological replicates; *p*-values calculated by two-sided Student’s t-test. K. Representative images of primary cultures of Sca-1+ and Sca-1- *Lgr5*-Apc^**f/f**^-tdT organoids. L. Schematic (left), quantification (middle), and representative images (right) of secondary cultures of Sca-1+ and Sca-1- *Lgr5*-Apc^**f/f**^-tdT organoids. Data expressed as mean ± S.D (n = 5); *P*-value calculated by two-sided Student’s t-test.

**fig. S5.**
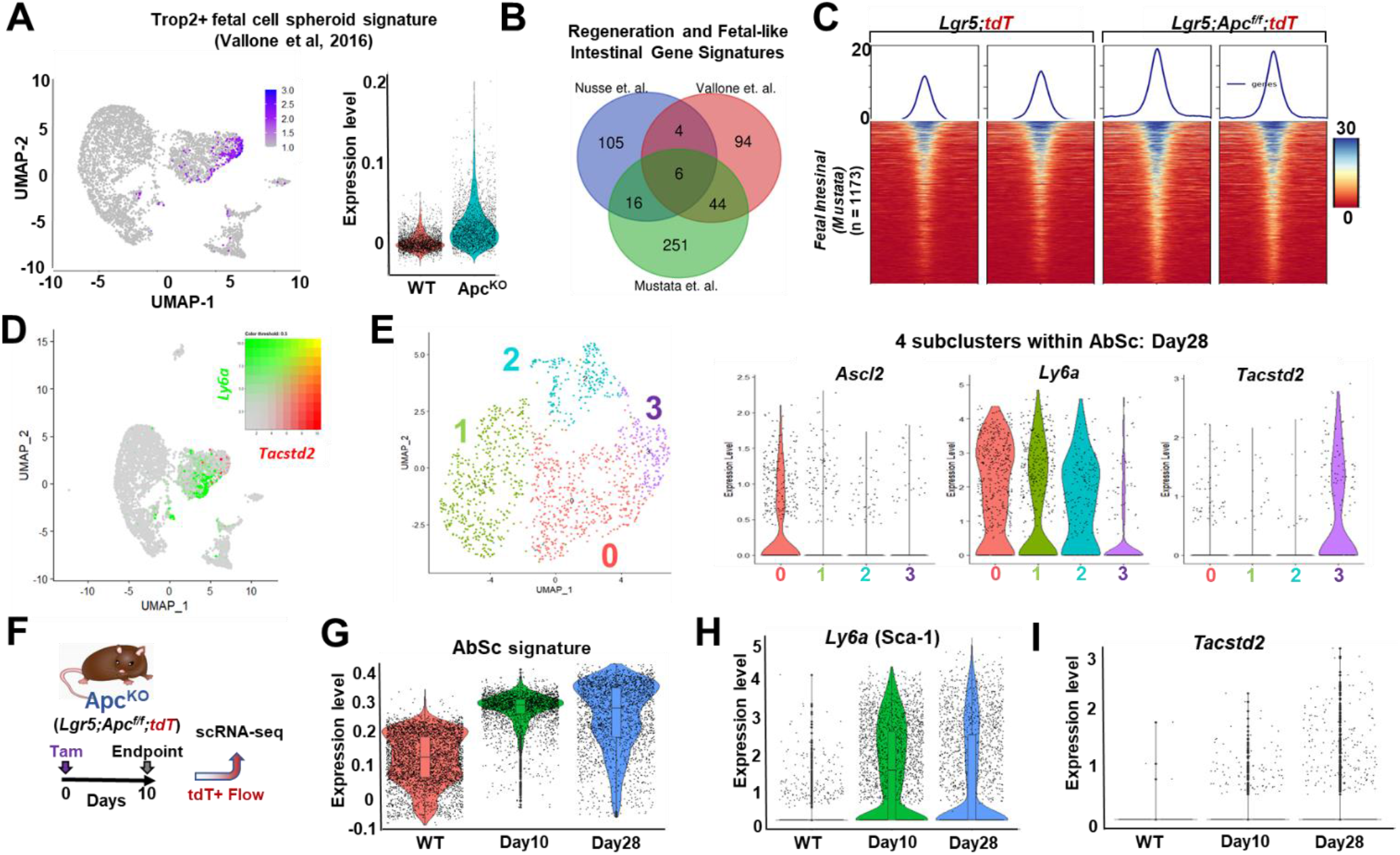
Regeneration precedes developmental reprogramming in AbSC cells from APC^KO^ mouse model. A. UMAP representation (left) and Violin plot (right) of experiment in Fig. 1F colored by Trop2+ fetal cell GI signature (Vallone et al.2016). B. Venn diagram showing the overlap of genes within the fetal-like intestinal and regenerative gene signatures C. Heatmap depicting chromatin accessibility at fetal genes in Lgr5;tdT and Lgr5;Apc^f/f^;tdT intestines using published signature (Mustata et al.) D. UMAP representation and color threshold (inset) of *Tacstd2* and *Ly6a* expression of experiment in Fig. 2A. E. UMAP representation of high-resolution sub clustering in AbSc cells from in Fig. 1F (left) and Violin plot of stem (*Ascl2*), regeneration (*Ly6a*) and fetal (*Tacstd2*) genes in the four subclusters (right). F. tdT+ intestinal epithelial cells isolated by fluorescence-activated cell sorting (FACS) from Lgr5;Apc^f/+^;tdT (control) and Lgr5;Apc^f/f^;tdT (experimental) mice 10 days following tamoxifen induction. G. Violin plot indicated expression of AbSC gene signature expression (from Day 28) in two different Apc^KO^ timepoints (Day10 and Day28) compared with Apc^WT^. H. Violin plot indicated expression of regeneration gene (*Ly6a*) in two different Apc^KO^ timepoints (Day10 and Day28) compared with Apc^WT^. I. Violin plot indicated expression of fetal gene (*Tacstd2*) in two different Apc^KO^ timepoints (Day10 and Day28) compared with Apc^WT^.

**fig. S6.**
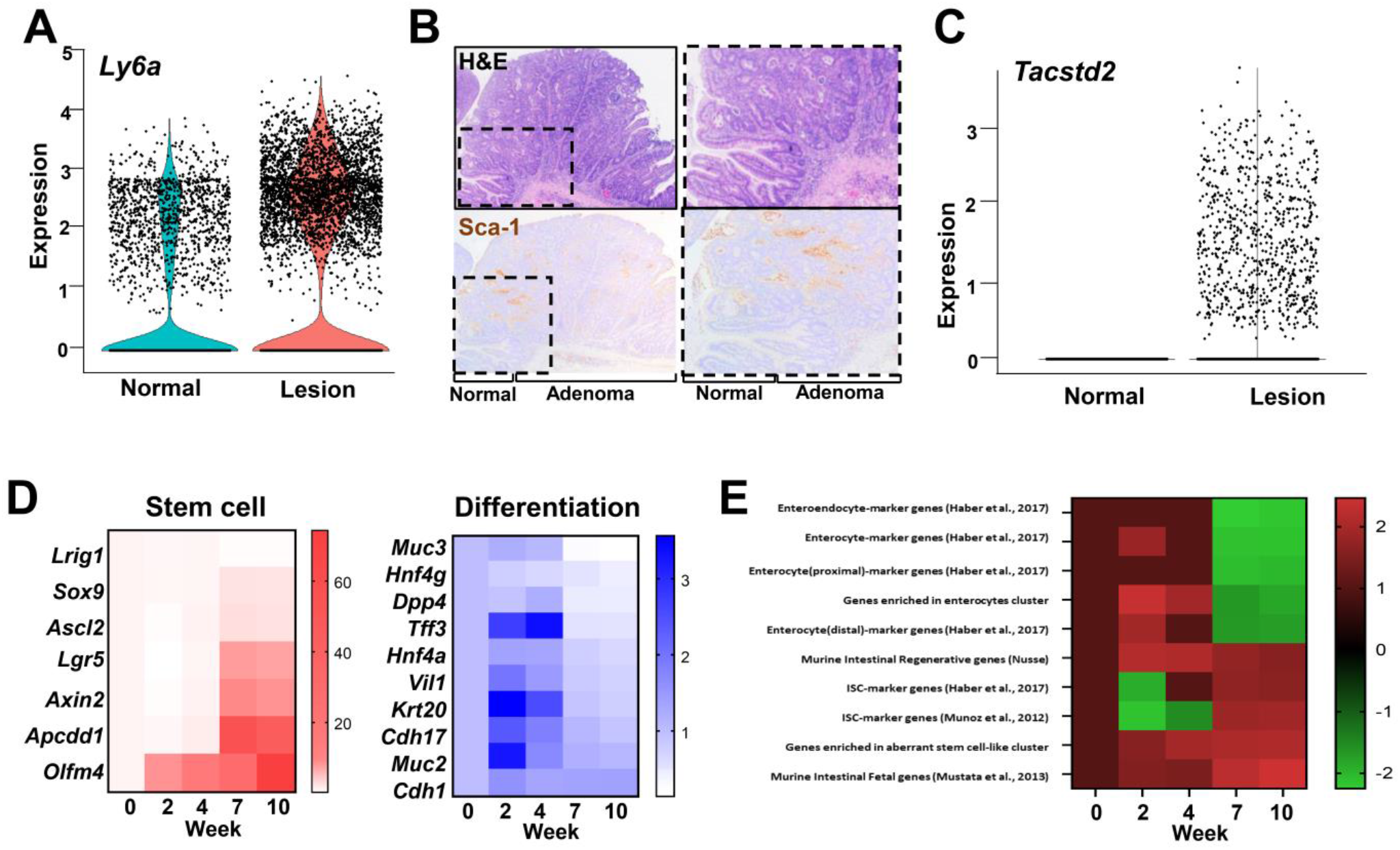
Regeneration precedes developmental reprogramming in AOM/DSS mouse model. A. Violin plot indicated expression of regeneration gene (*Ly6a*) in normal and neoplastic tissue from AOM/DSS mouse model scRNA-seq data. B. Representative images of H&E staining and Sca-1 from normal tissue and high-grade dysplastic lesion from experiment in Fig. 1C and Fig. 3A. C. Violin plot indicated expression of fetal gene (*Tacstd2*) in normal and neoplastic tissue from AOM/DSS mouse model scRNA-seq data. D. Heatmap depicting expression of Stem cell and differentiation genes for tumors at Week 10 vs controls from Chen et.al, 2022. E. Heatmap depicting normalized enrichments scores for different relevant signatures over time in AOM/DSS data from Chen et.al, 2022.

**fig. S7.**
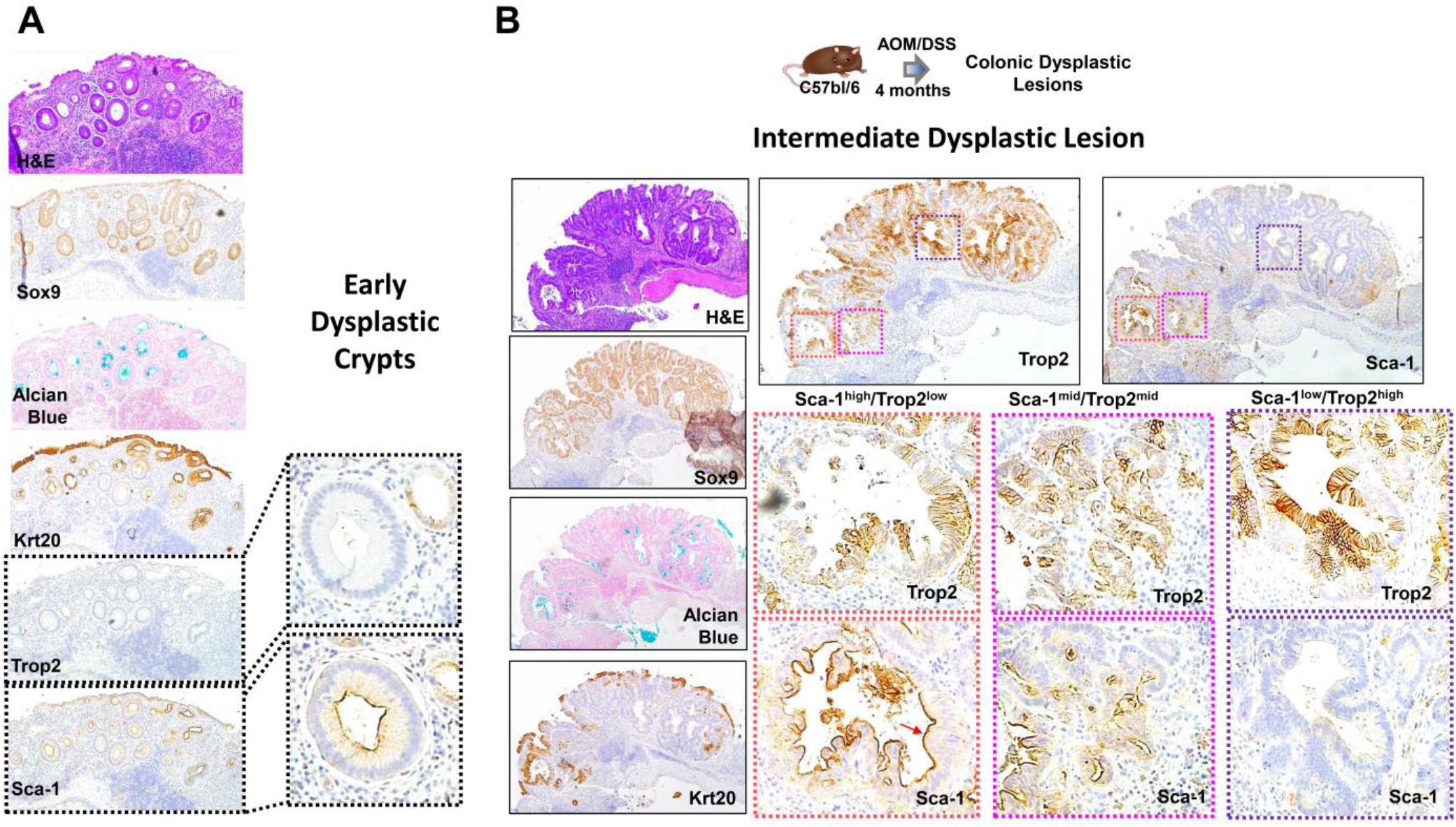
Sca-1 and Trop2 staining in early and intermediate dysplastic lesions from AOM/DSS mouse model. A. Representative images of H&E and Alcian Blue staining as well as Sox9, Krt20, Trop2, and Sca-1 from dysplastic colon crypts from an AOM/DSS treated mouse. B. Representative images of H&E and Alcian Blue staining as well as Sox9, Krt20, Trop2, and Sca-1 from dysplastic colon crypts from an AOM/DSS treated mouse.

**fig. S8.**
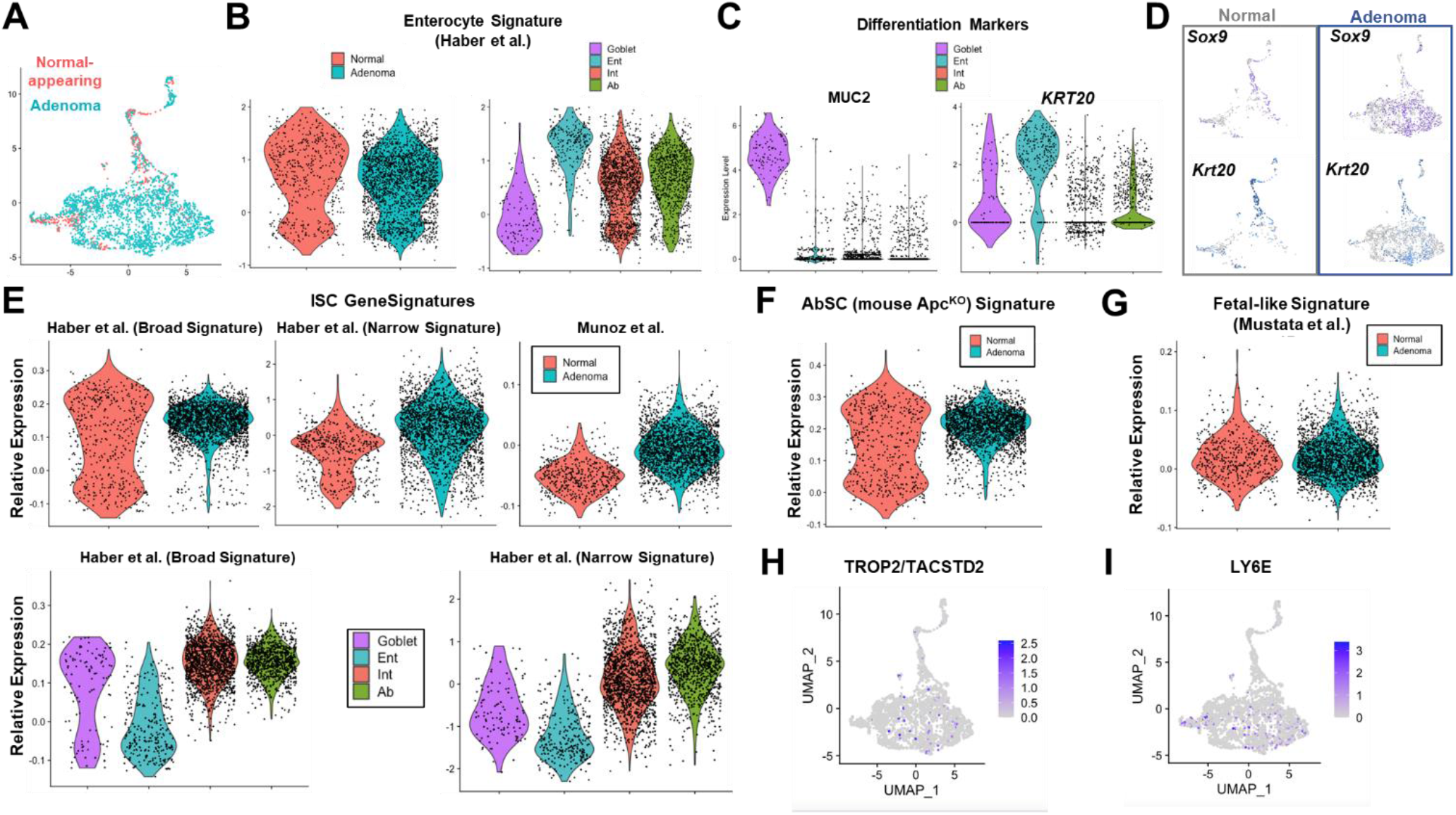
Genes and pathways in human FAP adenoma by scRNA-seq. A. UMAP representation of scRNA-seq of adenoma (blue) and paired normal (red) tissue from a patient with FAP. B. Violin plot indicating normalized expression of enterocyte signature (Haber et. al.) in human adenoma and paired normal (left) and in different cell clusters (right); Ent = Enterocyte, Int = Intermediate, and Ab = aberrant stem cell-like. C. Violin plot indicating normalized expression of differentiation markers MUC2 (left) and KRT20 (right) in different clusters. D. Normalized expression of *SOX9* and *KRT20* on UMAP plot separated by sample E. Violin plot indicating normalized expression of ISC gene signatures from (Haber et. al.) and (Munoz et. al.) shown by sample and cluster type. F. Violin plot indicating normalized expression of AbSC gene signature by sample type. G. Violin plot indicating normalized expression of fetal-like gene signature (Mustata et al.) by sample type. H. Normalized expression of *TROP2/TACSTD2* on UMAP plot. I. Normalized expression of *LY6E* on UMAP plot.

**fig. S9.**
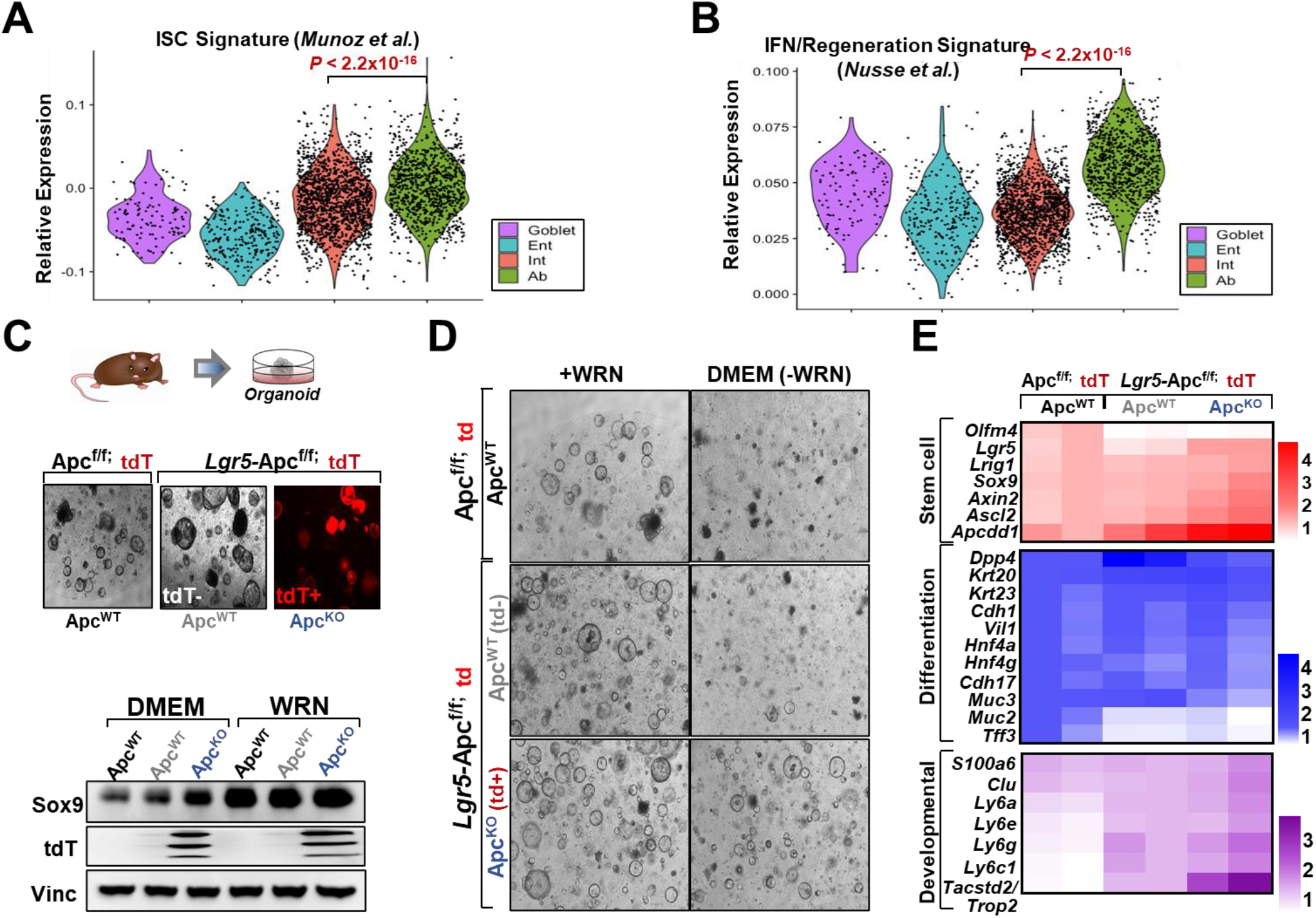
Genes and pathways in human FAP by scRNA-seq. A. Violin plot indicated expression of ISC (Munoz et al.) gene signature in four different cell clusters. P-value calculated by Wilcoxon rank sum test with Bonferroni correction. B. Violin plot indicated expression of IFN/fetal-like gene signature (Nusse et al.) in four different cell clusters. P-value calculated by Wilcoxon rank sum test with Bonferroni correction. C. Phase contrast and fluorescent images of intestinal organoids derived from Apc^f/f^;tdT (control) and Lgr5;Apc^f/f^;tdT mice (top). Immunoblot of Sox9, tdT, and Vinculin (loading control) protein levels in indicated organoids (bottom). D. Phase contrast and fluorescent images of intestinal organoids derived from Apc^f/f^;tdT (control) and Lgr5;Apc^f/f^;tdT mice grown in WRN and DMEM media. E. Expression of stem cell (red), differentiation (blue) and developmental (purple) genes in organoids derived from Apc^f/f^;tdT (control) and Lgr5;Apc^f/f^;tdT mice by bulk RNA-sequencing.

**fig. S10.**
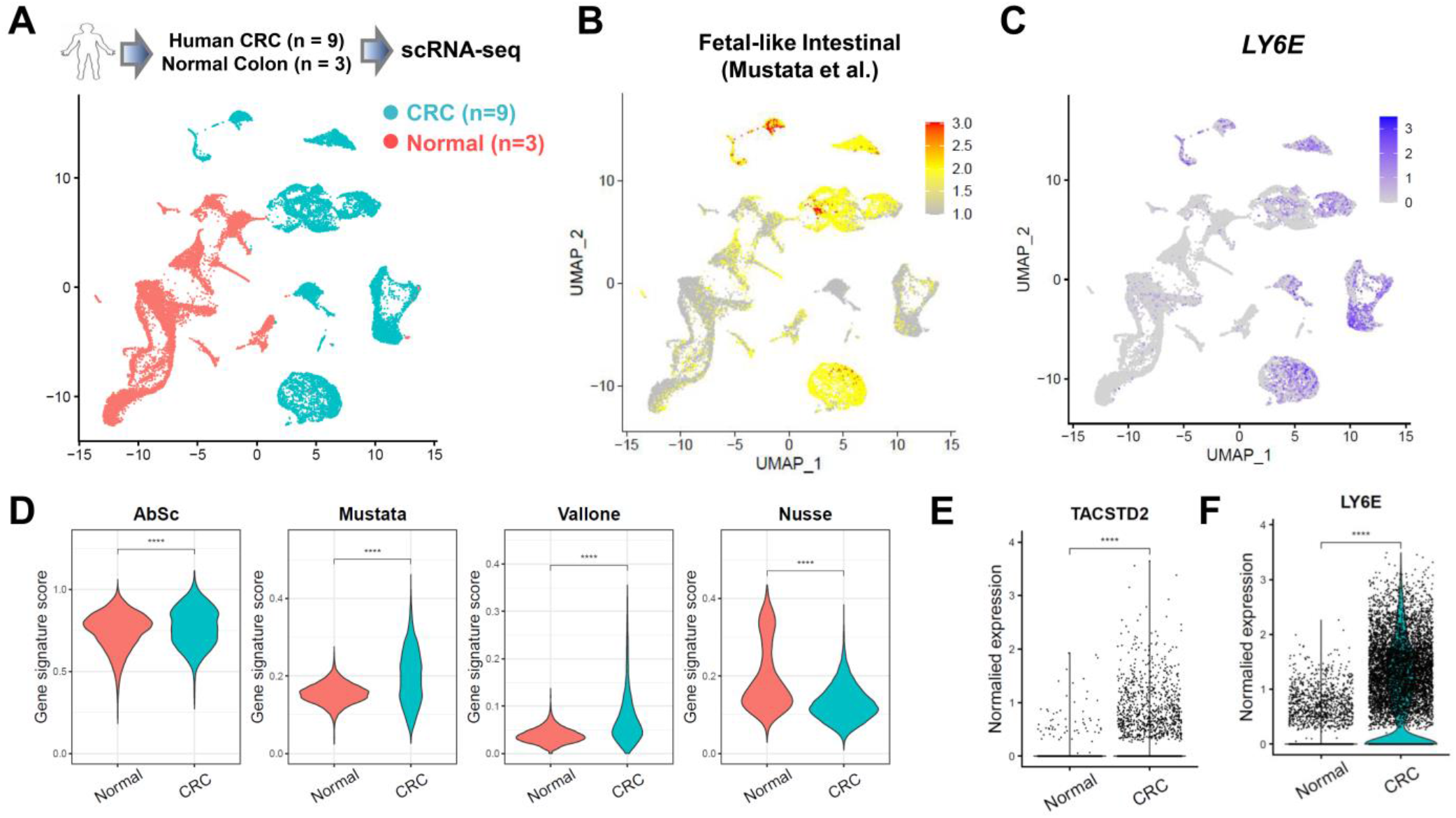
Genes and pathways in human CRC by scRNA-seq. A. UMAP representation of scRNA-seq of normal mucosa from healthy donors (pink, n=3) and intestinal tissue from patients with CRC (blue, n=9). B. UMAP representation of fetal-like gene signature (Mustata et al.) on UMAP plot. C. UMAP representation of *LY6E*. D. Violin plots of AbSC, Mustata, Vallone, and Nusse gene signatures in normal colon and CRC scRNA-seq data E. Violin plot of *TACSTD2* expression in normal colon and CRC scRNA-seq data. F. Violin plot of *LY6E* expression in normal colon and CRC scRNA-seq data.

**fig. S11.**
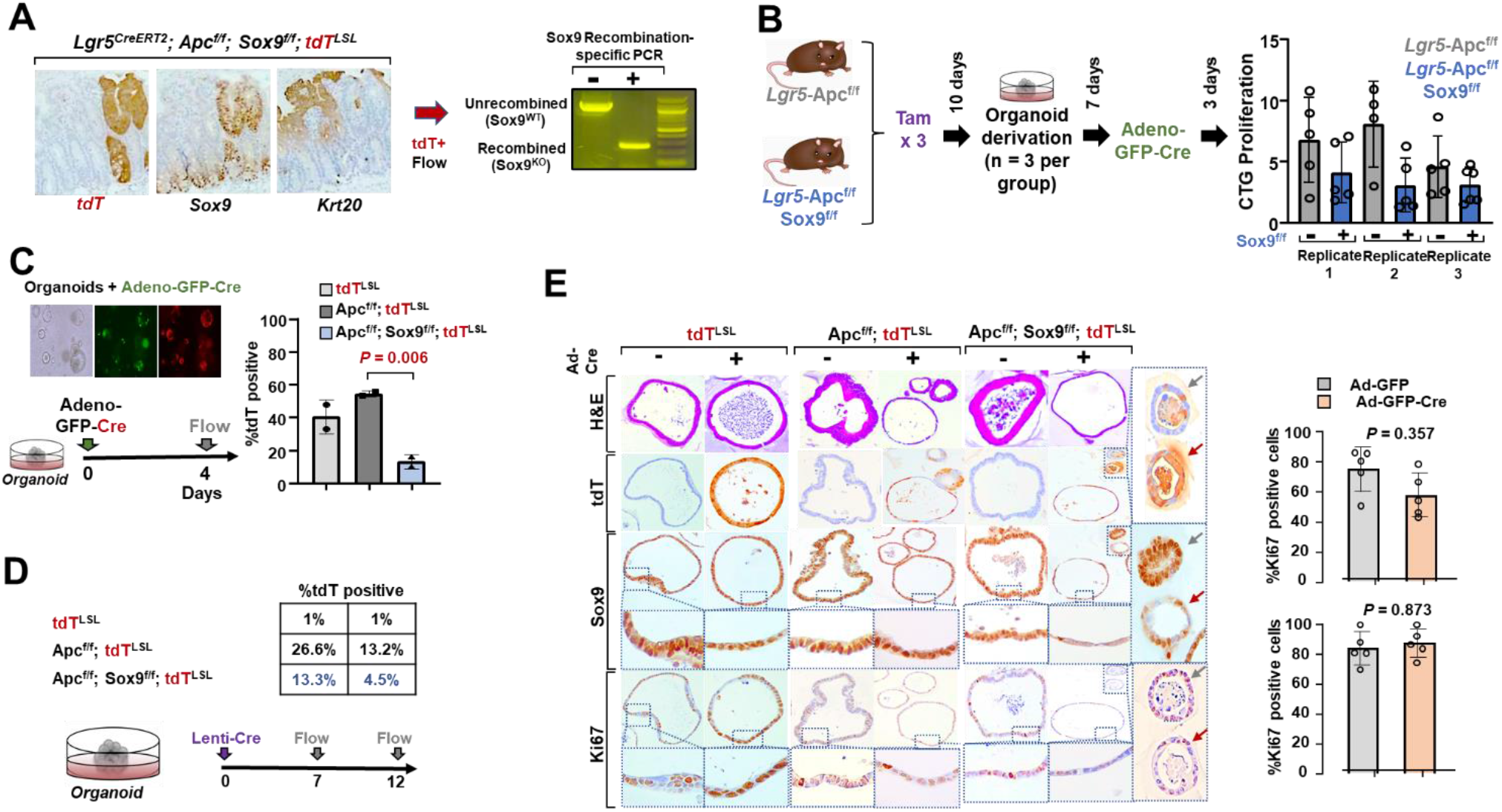
Sox9 is required for Apc^KO^ adenomas and organoids. F. Representative images of tdT, Sox9, and Krt20 IHC from *Lgr5*-Apc^f/f^-Sox9^f/f^-tdT mice. Genomic DNA was extracted from tdT+ cells isolated by FACS and subjected to *Sox9* recombination-specific PCR. PCR product indicating no recombination ~ 1200bp and recombination ~ 300bp. G. Schematic (left) and analysis (right) of an *in vivo* experiment of *Lgr5*-Apc^f/f^-tdT and *Lgr5*-Apc^f/f^- Sox9^f/f^-tdT mice in which organoid were generated 10 days following tamoxifen injection and then treated with Ad^GFP-Cre^ and subjected to a proliferation assay by CTG. Data expressed as mean ± S.D of five biological replicates from three mice per genotype. H. Organoids derived from tdT, Apc^f/f^-tdT, and Apc^f/f^-Sox9^f/f^-tdT mice were infected with Ad^GFP-Cre^ and percent tdT+ was quantified by FACS using the experimental approach in the schematic. Data expressed as mean ± S.D of two independent experiments. *P*-values calculated by two-sided Student’s t-test I. Organoids derived from tdT, Apc^f/f^-tdT, and Apc^f/f^-Sox9^f/f^-tdT mice were infected with Lenti^Cre^ and percent tdT+ cells were quantified by FACS using the experimental approach in the schematic. J. Organoids derived from tdT, Apc^f/f^-tdT, and Apc^f/f^-Sox9^f/f^-tdT mice were infected with Ad^GFP^ or Ad^GFP-Cre^, formalin fixed, and then processed for histopathology. Representative images of H&E staining and tdT, Sox9, and Ki67 IHC. Quantification of %Ki67 positivity in 5-6 organoids per condition. *P*-values calculated by two-sided Student’s t-test

**fig. S12.**
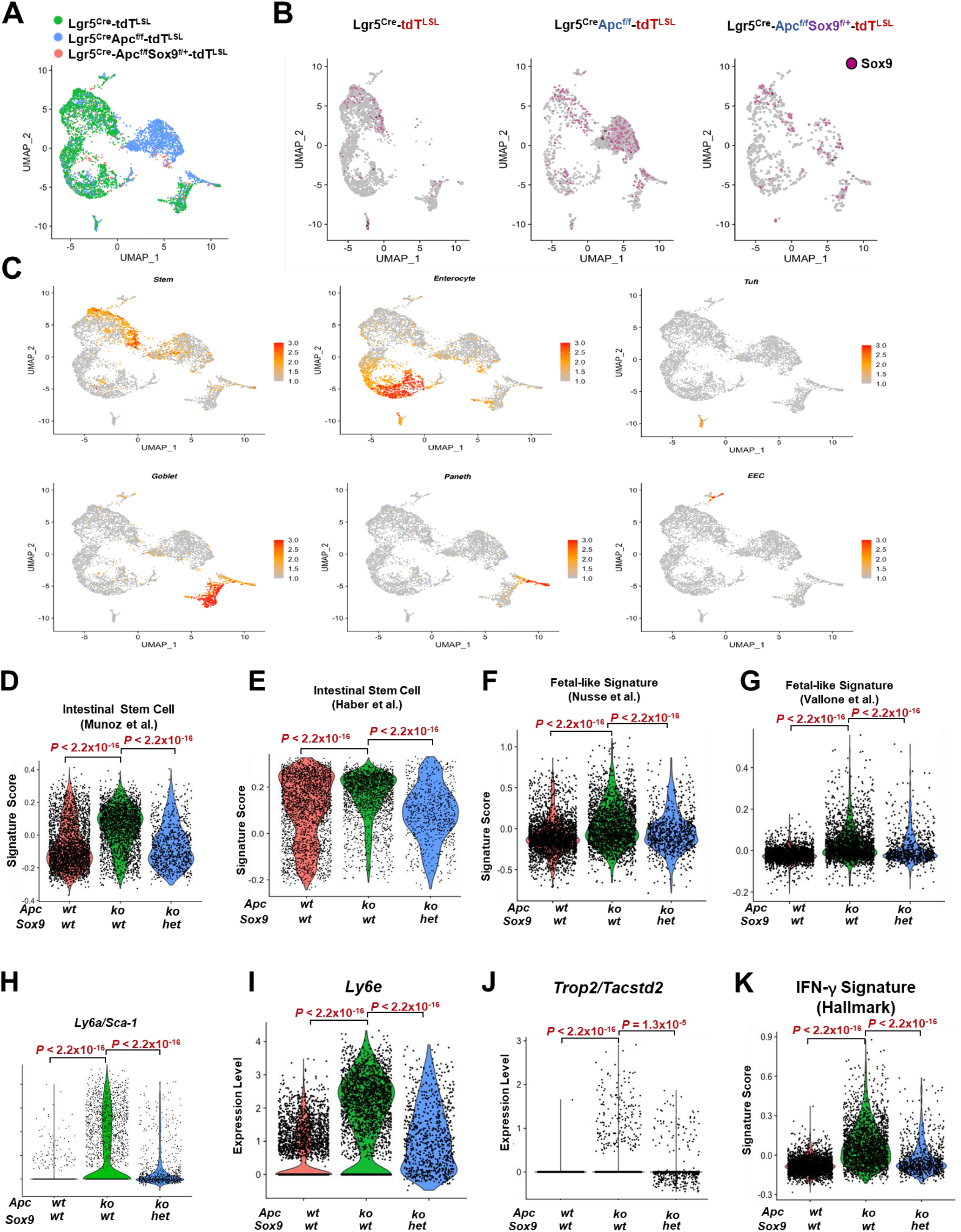
Sox9 attenuation leads to reduction in pathways associated with AbSC by scRNA-seq. A. UMAP projection of scRNA data of tdT+ cells from *Lgr5;tdT* (green), *Lgr5*;Apc^f/f^;tdT (blue), and *Lgr5*;Apc^f/f^;Sox9^f/f^;tdT (pink). B. Normalized expression of *Sox9* on UMAP plot separated by sample. C. Normalized expression of ISC, Enterocyte, Tuft, Goblet, Paneth, and Enteroendocrine (EEC) gene signatures. D. Violin plot showing single cell expression of ISC gene signature (Munoz et. al.) in samples. *P*-values calculated by Wilcoxon rank sum test with Bonferroni correction. E. Violin plot showing single cell expression of ISC gene signature (Haber et. al.) in samples. *P*-values calculated by Wilcoxon rank sum test with Bonferroni correction. F. Violin plot showing single cell expression of regeneration gene signature (Nusse et. al.). *P*-values calculated by Wilcoxon rank sum test with Bonferroni correction. G. Violin plot showing single cell expression of fetal-like gastrointestinal gene signature (Vallone et al.). *P*-values calculated by Wilcoxon rank sum test with Bonferroni correction. H. Violin plot showing single cell expression of *Ly6a. P*-values calculated by Wilcoxon rank sum test with Bonferroni correction. I. Violin plot showing single cell expression of *Ly6e. P*-values calculated by Wilcoxon rank sum test with Bonferroni correction. J. Violin plot showing single cell expression of *Tacstd2. P*-values calculated by Wilcoxon rank sum test with Bonferroni correction. K. Violin plot showing single cell expression of IFN-γ gene signature (Hallmark). *P*-value calculated by Wilcoxon rank sum test with Bonferroni correction.

**fig. S13.**
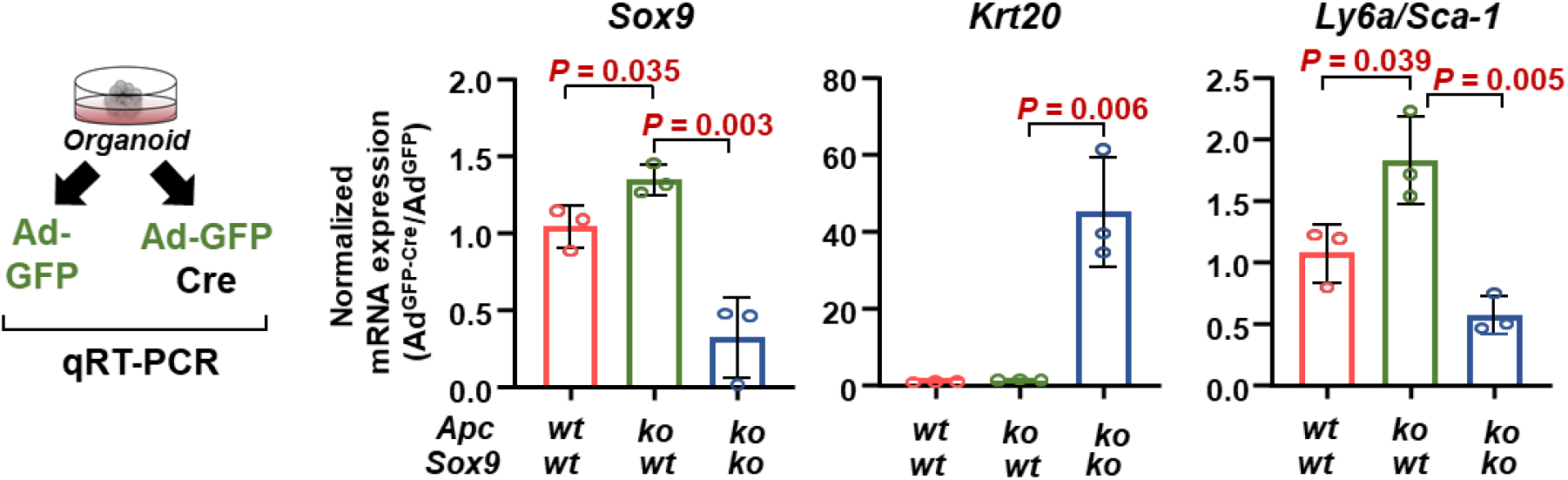
Sox9 attenuation leads to reduction in pathways associated with AbSC by qRT-PCR. mRNA expression of *Sox9, Krt20*, and *Ly6a* in organoids derived from tdT, Apc^f/f^;tdT, and Apc^f/f^;Sox9^f/f^;tdT mice expressed as a ratio of Ad^GFP-Cre^ to Ad^GFP^ treated (control). Data expressed as mean ± S.D of three biological replicates. P-values calculated by two-sided Student’s t-test.

**fig. S14.**
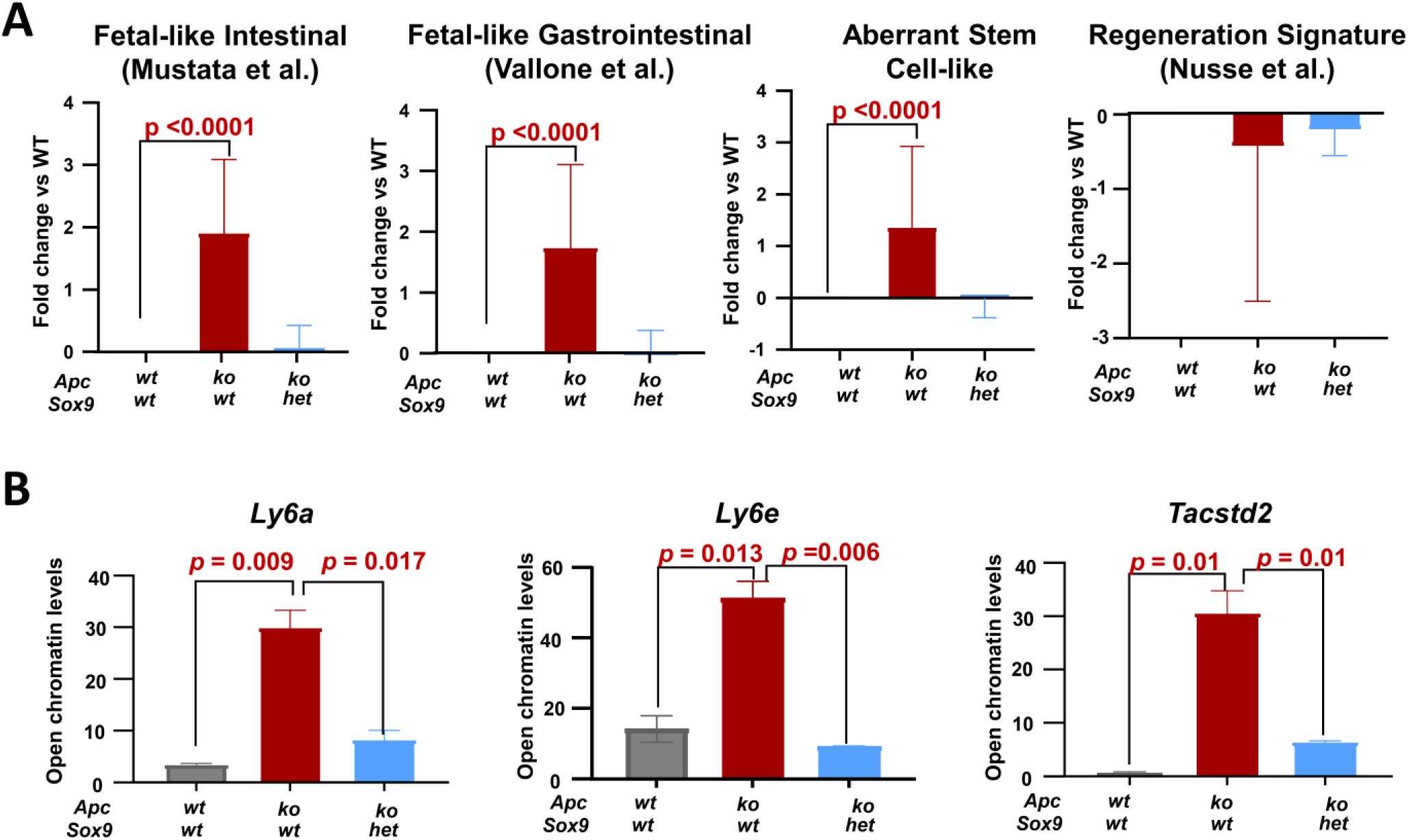
Quantification of chromatin accessibility changes in genetic mouse model. A. Quantification of chromatin accessibility in AbSC, fetal-like gastrointestinal (Mustata and Vallone) and regeneration (Nusse) genes derived from ATAC-sequencing of tdT+ cells from intestines of *Lgr5*;tdT, *Lgr5*;Apc^f/f^;tdT, and *Lgr5*;Apc^f/f^;Sox9^f/+^;tdT mice. B. Quantification of chromatin accessibility at *Ly6a, Ly6e* and *Tactsd2* genes derived from ATAC-sequencing of tdT+ cells from intestines of *Lgr5*;tdT, *Lgr5*;Apc^f/f^;tdT, and *Lgr5*;Apc^f/f^;Sox9^f/+^;tdT mice.

**fig. S15.**
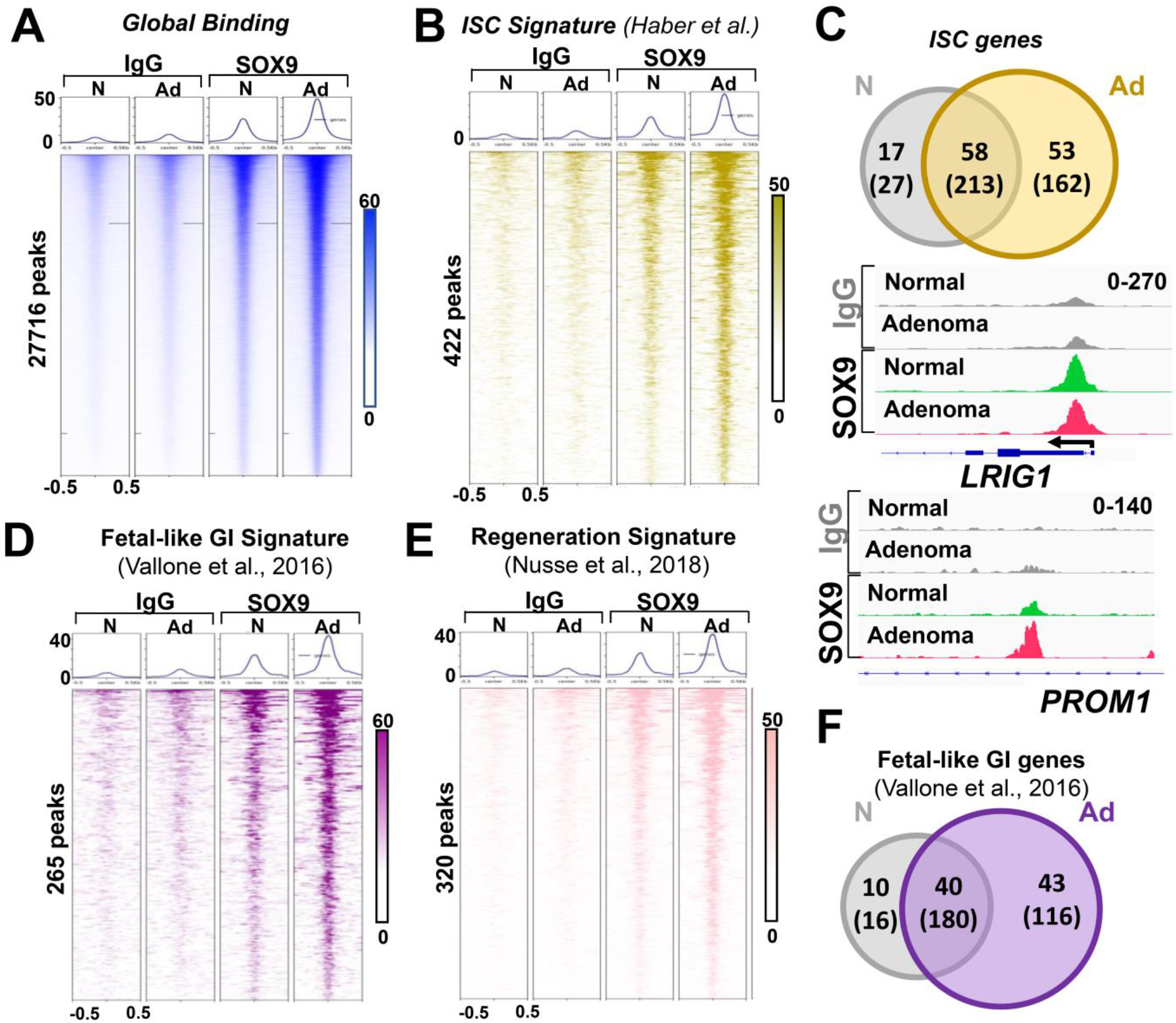
SOX9 binding in normal colon and adenoma FAP organoids by CUT&RUN. A. Heatmap of global SOX9 binding in normal colon and adenoma organoids from a patient with FAP by CUT&RUN B. Heatmap of SOX9 binding associated with ISC genes (Haber et al.) in normal colon and adenoma organoids from a patient with FAP by CUT&RUN C. Venn diagram of high-confidence SOX9 bound genes (peaks) using LFC ≥ 0.5 compared to IgG control in normal colon and adenoma organoids for ISC genes (top) and SOX9 binding at ISC gene *LRIG1* and AbSC gene *PROM1* in normal colon and adenoma organoids by CUT&RUN. D. Heatmap of SOX9 binding associated with fetal-like GI genes (Vallone et al.) in normal colon and adenoma organoids from a patient with FAP by CUT&RUN E. Heatmap of SOX9 binding associated with regeneration genes (Nusse et al.) in normal colon and adenoma organoids from a patient with FAP by CUT&RUN F. Venn diagram of high-confidence SOX9 bound genes (peaks) using LFC ≥ 0.5 compared to IgG control in normal colon and adenoma organoids for fetal-like GI genes

**fig. S16.**
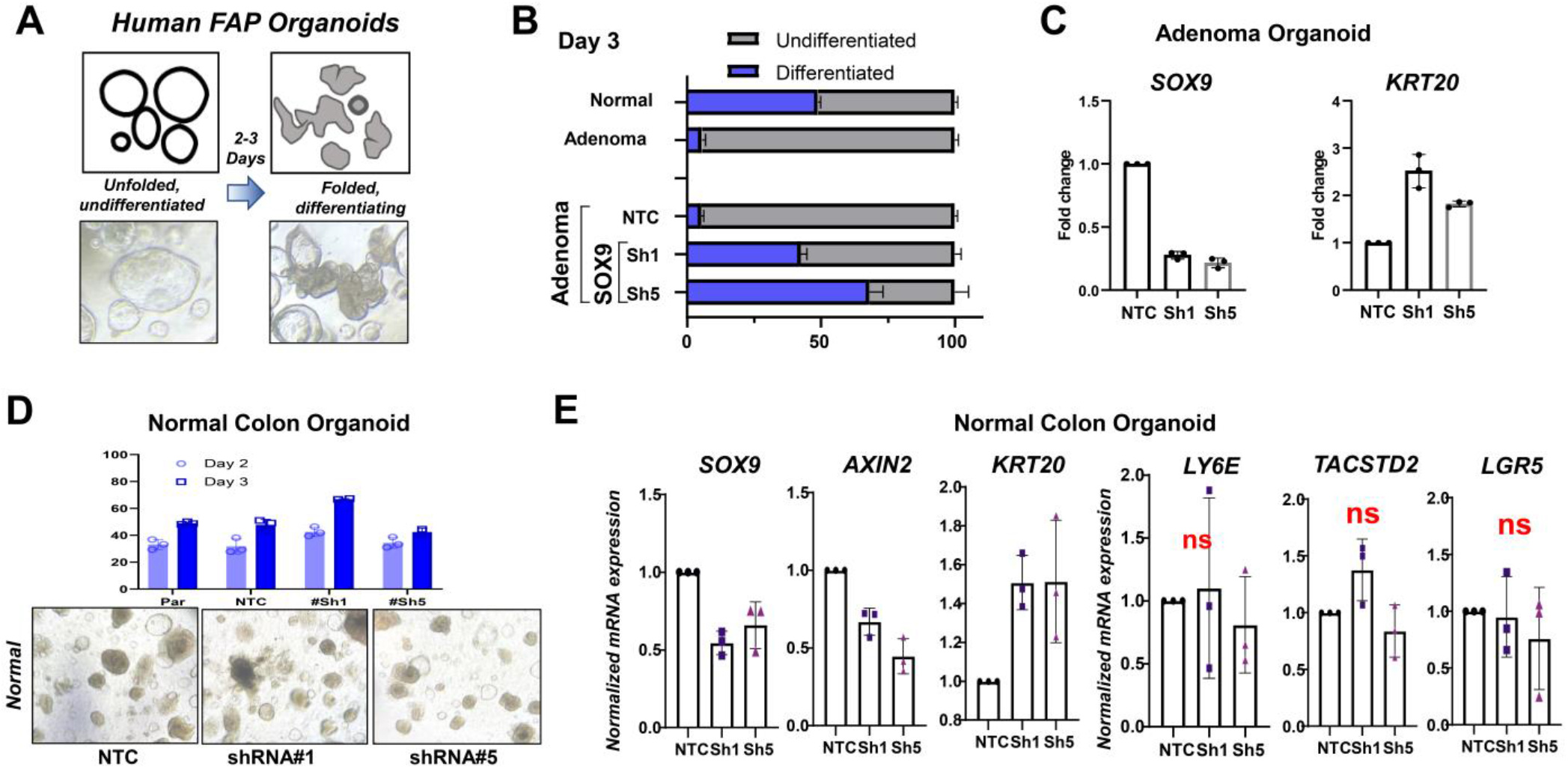
SOX9 necessity in normal colon and adenoma FAP organoids. A. Graphical representation (top) and phase contrast images (bottom) of differentiation phenotype in organoids. B. Quantification of percent undifferentiated and differentiated organoids in indicated cultures at day 3; mean ± S.D. of three biological replicates. C. Relative mRNA expression of *SOX9* and *KRT20* in FAP adenomas expressing non-targeting (NT) control, SOX9 shRNA#1, or SOX9 shRNA#5 by qRT-PCR; mean ± S.D. of three biological replicates. D. Quantification of percent undifferentiated and differentiated normal colon organoids expressing NTC or two independent shRNAs against SOX9 at day 2 and 3; mean ± S.D. of three biological replicates; representative images below. E. Relative mRNA expression of *SOX9, AXIN2, KRT20, LY6E, TACSTD2*, and *LGR5* in normal colon organoids expressing non-targeting (NT) control, SOX9 shRNA#1, or SOX9 shRNA#5 by qRT-PCR; mean ± S.D. of three biological replicates.

## METHODS

### Animal studies

All procedures involving mice and experimental protocols were approved by Institutional Animal Care and Use Committee (IACUC) of Dana-Farber Cancer Institute (11-009).

The generation of *Lgr5*^eGFP-IRES-CreERT2^ mouse was described earlier(*69*). *Lgr5*^eGFP-IRES-CreERT2^ mice were backcrossed in C57bL/6J mice and subsequently SNP tested to ensure >97% pure background (Taconic). To inactivate Apc in intestinal tissue, we crossed Apc^f/f^ mice (a generous gift from Christine Perret) to *Lgr5*^eGFP-IRES-CreERT2^. These mice were further crossed to *R26*-LSL-tdTomato purchased from Jackson Laboratory (JAX stock #007905) (*70*) for conditional tdT labeling of Lgr5+ stem cells and their progeny. To delete Sox9, mice were crossed to Sox9^flox^ mice (JAX stock #013106) (*53*).

To activate conditional alleles, experimental mice aged 6–8 weeks were injected intraperitoneally with a single dose tamoxifen (50 ul of a sunflower oil at 10 mg/ml) unless otherwise indicated. For MNU model, *Lgr5*^eGFP-IRES-CreERT2^ mice were treated with drinking water containing 240ppm of N-methyl-N-nitrosourea (MNU; biokemix) scheduled every other week for 10 weeks and followed for 1 year. For AOM/DSS model, AOM/DSS treatment was performed as previously described (*71*). Briefly, at 8-10 weeks of age mice were injected intraperitoneally with 10 mg/kg AOM (Sigma-Aldrich) and treated with 2% DSS (30-50 kDa, colitis grade, MP Biomedicals) in drinking water for 5 days. DSS treatment was repeated 3 times once per 4 weeks. Mice were sacrificed 11 weeks after AOM injection.

Mice were euthanized at the endpoint of the experiment and small intestine and colon were removed from mice, flushed with PBS, and incubated in 10% formalin for 5 min. Subsequently, intestines were opened longitudinally, ‘swiss-rolled’, incubated overnight in 10% formalin at room temperature and processed for paraffin embedding. Fresh tissue was collected for organoid generation, RNA isolation, protein collection and flash-frozen for long-term storage.

### Human specimen

Adenoma and paired normal-appearing tissue from a patient with familial adenomatous polyposis were collected post-colectomy under approval (protocol 13-189) by the Internal Review Board of the Dana Farber Cancer Institute, Boston, Massachusetts, USA.

### Histopathology

Paraffin-embedded intestines, organoids or xenograft tumors were serially sectioned and mounted on a slide. Sections were subjected to hematoxylin and eosin (H&E), Alcian blue-Periodic Acid Schiff (AB-PAS), as well as immunostaining, using standard procedures. For morphological analysis, sections were serially dehydrated in xylene and ethanol, stained with H&E for histological assessment or AB-PAS to identify goblet cells and mucus.

### Immunofluorescence and immunohistochemistry

For immunostaining, antigen retrieval was performed using a sodium citrate buffer (pH6), Trilogy (Sigma Aldrich Cell Marque) or Tris-EDTA pH9. Slides were permeabilized using a 0.2% Triton X100 for 30 minutes at room temperature and blocked with donkey serum for 1 hour. The primary antibodies used for immunohistochemistry were rabbit anti-Sox9 (1:600, CST, #82630,Trilogy antigen-retrieval), rabbit anti-Mucin2 (1:200, Santa Cruz, sc-15334, Tris-EDTA), anti-Krt20 (1:500, CST, #13063, Tris-EDTA), anti-RFP (1:500,Rockland, 600-401-379, sodium Citrate pH6), anti-eGFP (1:1000, Abcam, ab290, Tris-EDTA), anti-Olfm4 (1: 1000, CST, #39141, sodium Citrate pH6), anti-Lyz, anti-Sca-1/Ly6a (1:150, Abcam, ab109211, sodium Citrate pH6), anti-Axin2 (1:800, Abcam, ab32197, sodium Citrate pH6) and rabbit anti-Ki67 (1:1,000, CST, #12202, sodium Citrate pH6). Binding of primary antibody was detected with 3,39-diamino-benzidine-tetrahydrochloride-dihydrate and counterstained with hematoxylin.

IHC quantification was performed using Fiji Is Just Image J (FIJI) application on 10X images from Leica DM750 microscope. Ly6a or Sox9 positive cells were quantified and expressed as a percentage of total tdT+ cells in intestinal lesions. For organoids, the percentage of Ki67 positive cells were quantified and expressed as a percentage of total cells in an organoid.

### Flow analysis, sorting, and staining

Isolated intestines from euthanized mice were washed with ice-cold PBS; villi were scraped using glass slides and dissociated in 5mM EDTA in PBS at 4°C for 20 minutes, shaking every 5-7 minutes for about 30 seconds. The epithelial fraction was collected by centrifugation and incubated at 37°C for 30 minutes in pre-warmed 4x TrypLE. Single cells filtered through 70um filter post trypsinization, washed with complete media, and collected in FACS buffer.

For antibody staining, 100ul of resuspended cells in FACS buffer were incubated with 1ul APC labelled Sca-1 antibody (Thermofisher scientific, Catalog # 17-5981-82) for 20 mins on Ice, washed with PBS and collected in FACS buffer. Cells were resuspended in 1-2mL FACS buffer with DAPI and passed through another 70-micron filter before transferring to 40 micron filtered FACS tubes and sorted. Sorted cells were collected in 5mL of 50%FBS+50%DMEM complete with 10 uM Y27632 ROCK inhibitor.

### Single cell RNA sequencing

Murine intestines and human adenoma samples were processed as described above and then subjected to single cell RNA sequencing (scRNAseq) with or without cell hashing. If hashed, the mouse cells were stained with TotalSeq™-30301 Hashtag 1 (BioLegend #155861) and TotalSeq™-30302 Hashtag 2 (BioLegend #155863) antibodies; no hashing was used for human samples. Viable cells were washed and resuspended in PBS with 0.04% BSA at a cell concentration of 1000 cells/μL. About 17,000 viable mouse cells were loaded onto a 10× Genomics Chromium™ instrument (10× Genomics) according to the manufacturer’s recommendations. The scRNAseq libraries were processed using Chromium™ single cell 5’ library & gel bead kit (10× Genomics). Matched cell hashing libraries were prepared using single cell 5’ feature barcode library kit. Quality controls for amplified cDNA libraries, cell hashing libraries, and final sequencing libraries were performed using Bioanalyzer High Sensitivity DNA Kit (Agilent). The sequencing libraries for scRNAseq and scTCRseq were normalized to 4nM concentration and pooled using a volume ratio of 4:1. The pooled sequencing libraries were sequenced on Illumina NovaSeq S4 300 cycle platform. The sequencing parameters were: Read 1 of 150bp, Read 2 of 150bp and Index 1 of 8bp. The sequencing data were demultiplexed and aligned to mm10-3.0.0 using cell ranger version 3.1.0 pipeline (10× Genomics).

### Single cell RNA sequencing analysis

#### Pre-processing, alignment and gene counts

De-multiplexing, alignment to the transcriptome, and unique molecular identifier (UMI) collapsing were performed using the Cellranger toolkit provided by 10X Genomics.

#### General Clustering

Standard procedures for QC filtering, data scaling and normalization, detection of highly variable genes, and hashtag oligo (HTO) demultiplexing were followed using the Seurat v3 in RStudio. Cells with unique feature counts lower than 100 and greater than 25,000 as well as cells with greater than 25% mitochondrial DNA were excluded. Counts were log-normalized and scaled by a factor of 10,000 according to the default parameters when using the Seurat LogNormalize function. Variable features were identified, and the data were scaled using the default parameters (Ngenes = 2000) of the FindVariableFeatures FIG. caleData Seurat functions, respectively. HTOs were demultiplexed using the HTODemux function, and cells were identified as containing HTO-1 or HTO-2 based on their maximal HTO-ID signal. The cell population was filtered to contain only HTO-positive, singlet cells for further analysis. Principle component analysis (PCA) was completed on the remaining cells and 10 principal components were selected for clustering, tSNE, and UMAP analyses. Cells were visualized primarily using UMAP non-linear dimensional reduction (dims 1:10, resolution = 0.3), from which feature plots were generated to demonstrate distribution of gene expression and APC^WT^ versus APC^KO^ cells and expression levels of various marker genes throughout the population. Marker genes for each resulting cluster were found using the FindMarkers function with the minimum prevalence set to 25%. One cluster was determined to be immune cells based off of top marker genes and excluded from further analysis. Cluster identities were defined using known marker genes for intestinal epithelial cell types. For AOM/DSS model and human FAP samples, immune, stromal, and other supporting cell types were removed to focus the analysis on epithelial cells.

#### General analysis

scRNA-seq IntegrateData function in Seuratv4 was used to counteract batch effects among human tissue (Paired Normal A, Paired Normal B, Adenoma) and mouse model samples. Principal Component Analysis (PCA) was then completed on the integrated object and the quantity of principal components selected for clustering was determined using the integrated object’s elbow plot. Cells were then visualized primarily using UMAP non-linear dimensional reduction from which feature and violin plots were generated to demonstrate distribution of gene expression and expression levels of various marker genes and gene signatures throughout the population.

#### scRNA-seq gene signature analysis

To analyze existing gene signatures on our scRNA-seq data, the Seurat AddModuleScore function in Seurat v4 was used to calculate the average normalized and scaled gene expression of a given gene list in each individual cell. Specific cell types were identified using established marker genes and gene signatures (*38*). Gene signature scoring was then visualized with feature and violin plots. To generate novel gene signatures, the Seurat FindMarkers function was used to create lists of genes differentially expressed in one specified subset in comparison to another given subset. Minimum prevalence was set to 25%.

#### Pseudotime analysis

After Seurat processing, single-cell transcriptomic profiles of the cells belonging to the AbSC cluster were fitted with a principal graph using the learn_graph() function in Monocle3 (Cao, J. et. al., 2019). By assuming that the earliest cells would most resemble the wild-type intestinal stem cells, we ordered these cells along the learned graph whereby the earliest cells have the highest expression of ISC gene signatures.

### Assay for Transposase-Accessible Chromatin sequencing (ATAC-seq)

ATAC libraries were prepared as described previously(*72*). Briefly, flow-sorted tdT+ cells (25,000 cells in duplicates) were lysed to prepare nuclear pellets which then underwent transposition with TDE1 Enzyme (Illumina, 20034197). Tagmented DNA was purified using Zymo DNA Clean and Concentrator-5 Kit (cat# D4014) and the purified DNA was PCR amplified with NEBNext 2x MasterMix and Illumina adapters (table S4). The libraries were purified post-PCR using AMPure XP beads (Beckman Coulter). 150-bp paired-end reads were sequenced on a Novaseq instrument (Illumina).

### ATAC data processing

ATAC sequencing data was analyzed as described previously (PMID: 28648363). In brief, reads were mapped to the mm10/GRCm38 genome using Bowtie2 aligner version 2.3.5 (*72*). MACS2 was used to call peaks using the parameters ‘-t $bamfile -f BAMPE -n qc/macs/$base.macs2 -q 0.01 -g mm’ (*73*). Bigwig files were generated using deepTools bamCoverage with the options ‘bamCoverage --binSize 10 --smoothLength 30 -p 4 -- normalizeUsing RPGC --effectiveGenomeSize 2730871774 --extendReads $fragLength -b $file -of bigwig’ (PMID: 24799436). All bigwig and bed files were filtered using the ENCODE Blacklist. Only peaks with P-value < 0.00001 were considered for further analyses. Proximal peaks (-2kb to 2kb relative to TSS) called by macs2 were linked/annotated to genes by ChIPseeker version 1.34.1(74). Then, computeMatrix and plotHeatmap functions from deepTools(75) were used to visualize peaks.

### Cleavage Under Targets & Release Under Nuclease (CUT&RUN) sequencing

CUT&RUN for SOX9 and IgG control in Adenoma and Normal organoids was done using CUTANA ChIC/CUT&RUN kit (Epicypher #14-1048) following manufacturer’s protocol. Briefly, 5 × 10^5^ cells were captured with Concanavalin A, permeabilized using 0.01% digitonin and incubated with 0.5μg antibody (anti-Sox9 (CST, #82630) and IgG (# 13-0042; Epicypher) in 50μL antibody buffer (20mM HEPES at pH 7.5, 150mM NaCl, 0.5mM Spermidine, 1x protease inhibitor cocktails (EDTA-free tablet; Roche), 2 mM EDTA, 0.01% digitonin) overnight. After removing unbound antibody, pAG-MNase (20X) in 50μL cell permeabilization buffer was added to the cells and incubated for 10 min at RT. Then CaCl2 (2mM) was added to activate MNase and incubated at 4 °C for 2 hr. The reaction was stopped using 33μL of 2X STOP buffer and E. coli spike-in DNA (0.5ng) was added as a control. The DNA from the released chromatin in the supernatant was purified and then quantified using Qubit dsDNA HS assay kit (Agilent Technologies). CUT&RUN libraries were constructed using NEBNext Ultra II DNA library preparation kit as described previously (Skene et al., 2018) with a few modifications. Briefly, end repair and dA-tailing were conducted on 6 ng of CUT&RUN eluted DNA for 30 min at 20°C followed by 1hr at 50°C. After adaptor ligation for 30 min at 20°C, the DNA fragments were purified by 1.75X vol of AMPure XP beads (Beckman Coulter) followed by 10-12 cycles of PCR amplification with Next Ultra II Q5 master mix. The PCR products were purified with 1X vol of AMPure XP beads. After quantitative and qualitative analysis, libraries with different indexes were pooled and sequenced on Illumina Novaseq platform with paired-end 150-bp reads.

### CUT&RUN data processing

CUT&RUN sequencing data was analyzed as described previously(*73*), following the standard pipeline (https://yezhengstat.github.io/CUTTag_tutorial/index.html). In brief, paired-end 150-bp reads were aligned to GRCh38 human genome using Bowtie2 version 2.2.5(*74*) with options: --local --very-sensitive --no-mixed --no-discordant --phred33 -I 10 -X 700. Technical replicates (n = 4 per condition) were merged before reads alignment to increase the power of peak calling. macs2(*75*) was used to call peaks from bam files. For SOX9 CUT&RUN peak calling, parameters —t input_file –p 1e-5 –f BAM –keep-dup all –n out_name was used to call narrow peaks. To check the SOX9 binding profile and enhancer activity of specific gene sets, proximal peaks (-2kb to 2kb relative to TSS) called by macs2 were linked/annotated to genes by ChIPseeker version 1.34.1(*76*). Then, computeMatrix and plotHeatmap functions from deepTools(*77*) were used to visualize peaks.

### Intestinal Organoid Culture

Colonic glands were isolated, treated with EDTA, and then resuspended in 30-50 ul of Matrigel (BD Bioscience) and plated in 24-well plates. WNT/R-spondin/Noggin (WRN) containing DMEM/F12 with HEPES (Sigma-Aldrich) containing 20% FBS, 1% penicillin/streptomycin and 50ng/ml recombinant mouse EGF (Life Technologies) was used for culturing Apc^KO^ colon organoids. For the first 2-3 days after seeding, the media was also supplemented with 10 mM ROCK inhibitor Y-27632 (Sigma Aldrich) and 10 mM SB431542 (Sigma Aldrich), an inhibitor for the transforming growth factor (TGF)-β type I receptor to avoid anoikis. For passage, colon organoids were dispersed by trypsin-EDTA and transferred to fresh Matrigel. Passage was performed every 3-4 days with a 1:3–1:5 split ratio. For human colon organoid culture, the previous media was supplemented with antibiotics 100 ug/ml Primocin (Invivogen), 100 ug/ml Normocin (Invivogen); serum-free supplements 1X B27 (Thermo Fisher (Gibco)), 1X N2 (Thermo Fisher (Gibco)); chemical supplements 10mM Nicotinamide (Sigma), 500mM N-acetylcysteine (Sigma), hormone 50mM [Leu15]-Gastrin (Sigma), growth factor 100ug/ml FGF10 (recombinant human) ( Thermo Fisher) and 500nM A-83-01 (Sigma), which is an inhibitor of the TGF-β Receptors ALK4, 5, and 7.

### Organoid fixation for FFPE

Confluent organoids in 6-well plate were fixed in 10% formalin at 4 °C on rocker overnight. The fixed organoids were washed with PBS, collected by centrifugation at 2000 rpm, 3 mins. The supernatant was carefully aspirated; cell pellets were resuspended in 50-80ul of 2% agar and then immediately kept on ice. The solidified agar with organoids was kept in a cassette and processed to make FFPE blocks.

### Cell Proliferation Assays

Cell viability was quantified by measuring cellular ATP content using the CellTiterGlo Cell Viability assay (Promega) according to the manufacturer’s instructions. All experiments were performed in triplicate in 96-well plates.

### SOX9 recombination PCR

Mouse tails were incubated at 95°C in 75μl of the cell lysis buffer (5 mL sterile ddH2O, 7 μl 50% NaOH, 7 μl 0.5M EDTA, pH 8.0) followed by 15 minutes at 4°C and neutralized with 75 μl of neutralization buffer (40 mM Tris-HCl, pH ~5). 2 μl of supernatant was used for PCR amplification using primers designed for WT SOX9; Unfloxed SOX9 and Floxed SOX9 (table S4) using Phire Tissue Direct PCR master mix (Thermo Fisher Scientific #F170S). PCR cycling conditions were initial denaturation 94°C – 120s; 10x touchdown cycles with the following three steps: 94°C - 10s, 65°C (0.5°C increment each cycle) – 10s, 72°C – 90s; 25x cycles with the following three steps: 94°C – 10s, 60°C– 10s, 72°C– 90s; the last annealing 72°C–180s. The PCR products were analysed on 1% agarose gel.

### Organoid Adenoviral Transduction

To transduce colonic organoids, 0.5 M organoids were suspended in 500ul of WRN media with 10 uM Y27632 ROCK inhibitor in a 24 well ultra-low attachment plate. Adenovirus (University of Iowa, Gene Transfer Vector Core, Stock: Ad3786) High viral titer equal to 1010 PFU/mL) carrying GFP or GFP-Cre (1ul) was added to the organoids and the sealed plate is centrifuged 600g x 1 hr at 32 C followed by incubation at 37 C for 5-6 hrs. Post-incubation, the transduced organoids were washed with complete media and resuspended in about 200 ul Matrigel and plated in nunclon delta plates with WRN conditioned media with 10 uM ROCK inhibitor.

### Generation of stable colon organoids

All genetically manipulated colon organoid lines were generated using the protocol described here (*78*). shRNAs against *SOX9* were cloned into PLKO.1, TET-PLKO and TET-Cellecta vectors. To generate lentiviruses, expression vectors were co-transfected into HEK293T cells with the lentiviral packaging constructs psPAX2 and pMD2.G (VSV-G) in a 1:1:1 ratio using X-tremeGENE9 DNA Transfection Reagent (Roche) according to the manufacturer’s instructions. Cell culture media was changed the following day and lentiviral supernatant was harvested 48 h and 72 h later and filtered through a 0.45 μm filter (Millipore). Lentiviruses were aliquoted and stored at - 80 °C until use.

To transduce colonic organoids, spheroids in one well (24-well plate) were trypsinized and a one-fourth to one-eighth volume of cell suspension was used for each infection. Cells were resuspended in 500 μl lentiviral supernatant with 8 μg/mL polybrene and 10 mM Y-27632, centrifuged at 600g 32 °C 1 hours, and incubated for 6 hours in cell culture incubator. The infected cells were suspended in 30-50 ul of Matrigel and cultured with Wnt/R-spondin-deprived medium containing 10 mM Y-27632 and 10 mM SB431542. Colon organoids were selected with 3 μg/ml puromycin at 24 hours post-infection.

### RNA isolation and qPCR

Total RNA was isolated using the RNeasy Mini Kit (Qiagen, Germantown, MD, USA) and cDNA was synthesized using the iScript™ Reverse Transcription Supermix for RT-qPCR (Bio-Rad) according to the manufacturer’s instructions. Gene-specific primers for SYBR Green real-time PCR were either obtained from previously published sequences or designed by PrimerBLAST (https://www.ncbi.nlm.nih.gov/tools/primer-blast/) and synthesized by Integrated DNA Technologies or ETON biosciences. Real-time PCR was performed and analyzed using CFX96 Real-Time PCR Detection System (Bio-Rad Laboratories, Inc., Hercules, CA) and using Power SYBR Green PCR Master Mix (Thermo Fisher Scientific) according to the manufacturer’s instructions. Relative mRNA expression was determined by normalizing to *GAPDH* expression, which served as an internal control. See table S4 for primers used for qPCR.

### Immunoblot, antibodies and inhibitors

Immunoblot analysis was performed as previously described (*79*). Briefly, cells were lysed in RIPA buffer supplemented with a protease inhibitor cocktail (Roche). Whole cell extracts were resolved by SDS-PAGE, transferred to PVDF membranes, and probed with indicated primary antibodies. Bound antibodies were detected with horseradish peroxidase (HRP)-conjugated secondary antibodies and chemiluminescent HRP substrate. The following primary antibodies were used for western blotting (all from Cell Signaling Technologies, Beverly, MA, USA, unless otherwise indicated): anti-SOX9 (#82630, 1:1,000), anti-Vinculin (#13901, 1:1,000), anti-RFP (Rockland, 600-401-379, 1:500).

### Statistical Analysis and reproducibility

Experiments were performed in triplicate. Data are represented as mean ± s.d unless indicated otherwise. For each experiment, either independent biological or technical replicates are as noted in the Fig. legends and were repeated with similar results. Statistical analysis was performed using Microsoft Office, Prism 7.0 (GraphPad), or RStudio statistical tools. Pairwise comparisons between groups (that is, experimental versus control) were performed using an unpaired two-tailed Student’s *t*-test or Kruskal–Wallis test as appropriate unless otherwise indicated. For all experiments, the variance between comparison groups was found to be equivalent. Animals were excluded from analysis if they were euthanized due to health reasons unrelated to tumor volume endpoint.

## Author contributions

P.B. and J.P.R. co-led the study. P.B., D.A., and G.N.D. performed mouse and organoid experiments, analyzed data, and wrote/revised the manuscript. P.B., J.P.R., C.M., G.N.D, and P.D. performed scRNA-seq analyses and wrote/revised the manuscript. P.B, D.A., and Z.L. analyzed the CUT&RUN data. X.L. generated the SOX9 hairpins. S.M.M. helped with AOM/DSS mouse model. J.H. and M.Y. provided samples for human analysis. W.C.H. revised the manuscript. N.S.S. designed and supervised the study, performed experiments, analyzed data, wrote/revised the manuscript.

## Acknowledgements

We thank Sethi lab members, Hahn lab members, particularly Jon So, Ramesh Shivdasani and members of his lab, particularly Pratik Singh, James Cleary, Harshabad Singh, James DeCaprio, Matthew Meyerson, William Sellars, and Adam Sperling for insightful discussions and/or review of the manuscript; Christine Perret for kindly sharing the Apc^flox/flox^ mice developed in her laboratory; Aniket Gad and Lay-Hong Ang for assistance with immunohistochemistry assays; Shuqiang Li and Kenneth Livak for assistance with scRNA-seq; Dana-Farber/Harvard Cancer Center for the use of the Specialized Histopathology Core, which provided histology and immunohistochemistry service; Harvard Digestive Disease Center and NIH grant P30DK034854 for core services, resources, technology, and expertise. Dana-Farber/Harvard Cancer Center is supported in part by an NCI Cancer Center Support Grant # NIH 5 P30 CA06516. J.P.R. is supported by NIH K00 CA212221. This work was funded by the Claudia Barr Award and Virtual Scholar Award from the Department of Defense (CA201084) to N.S.S.

Supplementary Information is available for this paper

Correspondence and requests for materials should be addressed to N.S.S.

